# An injectable chitosan hydrogel localizes and tunably releases immunotherapeutics intratumorally eliminating both treated and abscopal murine triple negative breast tumors

**DOI:** 10.1101/2024.12.19.629422

**Authors:** Siena M Mantooth, Jarred M Green, William D Green, Khue G Nguyen, Kateri A Mantooth, Danielle M Meritet, J Justin Milner, David A Zaharoff

## Abstract

Systemic delivery of immunotherapy is dose-limited and often causes serious immune-related adverse events. Intratumoral injections can reduce systemic immunotoxicities and increase immunotherapy concentrations within a tumor. However, high pressures associated with direct tumor injection limits injectate retention, as low viscosity, saline-based solutions rapidly leak out of tumors. Viscoelastic solids, such as hydrogels, can improve local retention of co-formulated immunotherapies and provide sustained delivery. Prior work demonstrated that a chitosan-based hydrogel, XCSgel, was shear-thinning, self-healing, injectable, biocompatible, and clinically imageable. Here, we investigated XCSgel as a localized intratumoral delivery platform in the context of murine models of orthotopic triple-negative breast cancer. The intratumoral retention of immunotherapeutics co-formulated in XCSgel was characterized both *ex vivo* and *in vivo* via fluorescence imaging. Histopathological responses to intratumoral injections of XCSgel alone were scored by a veterinary pathologist. Initial antitumor studies evaluated a range of antitumor cytokines co-formulated with XCSgel. Subsequent antitumor and rechallenge studies evaluated the efficacy of a single intratumoral injection of interleukin-12 (IL-12) co-formulated in XCSgel (XCSgel-IL12) to control the growth of primary and abscopal tumors while inducing protective immunity. Pharmacokinetics studies quantified the systemic dissemination of IL-12 and consequent production of interferon-gamma following intratumoral injection with XCSgel co-formulation. Spectral flow cytometry was used to document changes in the tumor-immune microenvironment (TIME). XCSgel resisted tumor leakage and slowly released three model cytokines. XCSgel could be tuned for faster or slower release of embedded therapeutics. XCSgel-IL12 outperformed XCSgel formulations with other commonly used antitumor cytokines. A single injection of XCSgel-IL12 eliminated 86% E0771 and 20% mWnt orthotopic primary TNBC tumors. Mice rendered tumor-free resisted a live tumor challenge. XCSgel-IL12 also eliminated 67% untreated abscopal E0771 tumors. XCSgel-IL12 induced profound changes to the TIME, including a 3-fold reduction in the frequency of exhausted CD8^+^ T cells and a 3.2-fold increase in activated, proliferating CD8^+^ T cells. XCSgel is a promising localized delivery platform well-suited to enhance the retention and antitumor activity of potent immunotherapeutics. A single injection of XCSgel-IL12 can eliminate both primary and abscopal solid tumors, indicating that systemic immunotherapy may not be required for systemic control of cancer.

**Synopsis:** A novel injectable hydrogel, XCSgel, can localize and slowly release immunotherapies to eliminate primary and abscopal murine triple negative breast cancer tumors with a single injection.

## Introduction

Systemically administered immunotherapies are dose-limited and benefit only a minority of treated patients. Between 70-90% of patients treated with checkpoint inhibitors suffer from at least one immune-related adverse event (irAE), with grade 3+ irAEs occurring in 21% of patients (1,2). At a maximum tolerated dose, systemically administered immunotherapies, be they checkpoint antibodies or immune-stimulating cytokines, such as interleukin-2 (IL-2) or interferon-alpha (IFN-α), are unlikely to reach optimal concentrations within a tumor (3). Thus, irAEs limit the dose of current immunotherapies, which in turn limit clinical benefit. These dose-limitations also preclude the use of potent immunotherapies in earlier stage cancer patients who are likely to respond more favorably to immune-based interventions (4).

Direct intratumoral (i.t.) delivery can potentially reduce systemic irAEs while maximizing immunotherapeutic concentrations within tumors. Advances in image-guided injections have made practically all solid tumors accessible. Accordingly, interest in local injections has increased in recent years. As of May 31, 2024, 160 active and/or recruiting clinical trials involved i.t. delivery of an immunotherapeutic (clinicaltrials.gov, key search terms: “intratumoral” OR “intralesional” AND “cancer”). Publications from recently completed trials indicate that i.t. injections in numerous solid cancers are safe and feasible (5). Unfortunately, lower than expected clinical response rates have limited development in i.t. delivery of immunotherapeutics. To date, only one immunotherapy has been approved for i.t. or intralesional injection – T-VEC for advanced melanoma (5). A critical aspect that has been overlooked during the development of i.t. immunotherapies is local retention after injection (6,7). Low viscosity injectates are excluded from tumors due to high injection pressures, which either cause tumor cracking or rapid leakage via the needle track. Therefore, most immunotherapies injected i.t. in saline media are likely not retained within the tumor.

Unlike standard saline-based vehicles, hydrogels are viscoelastic solids that can resist shear forces and subsequent leakage from i.t. injections. Thus, immunotherapies co-formulated within hydrogels are more likely to be retained intratumorally following injection. Hydrogels may also be engineered for sustained, tunable release of therapeutics within a solid tumor. Numerous injectable and implantable hydrogels for immunotherapy delivery are in preclinical development (8). However, none have reached clinical trials.

To fill this gap, we recently developed a novel, injectable hydrogel, called XCSgel. Our previous *in vitro* characterization of this chitosan-based crosslinked hydrogel demonstrated that it is shear-thinning, self-healing, biocompatible, imageable and capable of providing sustained release of biologics (9). Chitosan is a naturally occurring polysaccharide, primarily derived from the shells of crustaceans and has been widely explored as a component of numerous delivery drug and gene delivery systems (10). In this study, we continue the characterization of XCSgel *in vivo* with a particular focus on treating triple-negative breast cancer (TNBC) via an XCSgel-localized immunotherapy.

Breast cancer (BC) diagnoses among women constituted about 2.3 million new cases in 2020, with 3 million projected in 2040 (11). Of the different subtypes of BC, TNBC has the lowest four-year survival rate (77%) and the highest likelihood of recurrence and metastasis (12). TNBC is more responsive to systemic immunotherapy, although response rates remain low around 20% (13). Therefore, we investigated the potential of localized immunotherapy, using XCSgel, to control TNBC and potentially improve the care of millions of patients worldwide.

The ability of XCSgel to resist expulsion following i.t. injection was characterized *in vitro* and *ex vivo*. Tissue responses to XCSgel delivered i.t. in an orthotopic TNBC murine model were characterized via histological analyses. *In vivo* retention and sustained release of three cytokines in XCSgel were demonstrated with fluorescence imaging. The antitumor efficacies of diverse antitumor cytokines were compared. Based on differences in efficacy, additional antitumor and pharmacokinetics studies of XCSgel when co-formulated with the cytokine IL-12 (XCSgel-IL12) were performed in multiple TNBC models. Changes in the tumor-immune microenvironment (TIME) following i.t. immunotherapy were evaluated with multi-spectral flow cytometry. Finally, XCSgel-IL12-induced systemic antitumor immunity was assessed via rechallenge studies and analyses of abscopal tumor control. Overall, studies suggest that XCSgel is a promising platform for the localization of immunotherapies within TNBC and likely other solid tumors.

## Materials and Methods

### Compounds, Reagents, and Animals

All compounds and reagents are listed in Supplemental Table 1. Recombinant murine interleukin-12 (IL-12) was overexpressed and purified in-house, as previously described (14). Seven- to 10-week-old female C57BL/6NCr and Balb/cAnNCr mice were purchased from Charles River Laboratories. Animal use complied with the Public Health Service Policy on Human Care and Use of Laboratory Animals. 203 total mice were used. All experiments involving laboratory animals were approved by the Institutional Animal Care and Use Committee at North Carolina State University (protocols 19-795, 22-414, and 23-126). ARRIVE reporting guidelines were used (15).

### XCSgel Preparation

XCSgel was prepared as previously described (9). XCSgel_slow_ refers to 95% deacetylated chitosan, at a molecular weight range 100-250kDa, crosslinked with 1:2 EDC:NHS molar ratio, at a concentration of 45 mg/mL. XCSgel_fast_ refers to 70% deacetylated chitosan, at a molecular weight range 100-250kDa, crosslinked with 1:2 EDC:NHS molar ratio, at a concentration of 30mg/mL, co-formulated with 3mg/mL carboxymethyl-chitosan (CM-CS). All XCSgels were prepared one day prior to delivery and stored overnight at 4°C.

### Cell Growth and Maintenance

E0771 cell lines were purchased from the American Type Culture Collection. The mWnt cell line was a kind and generous gift from Dr. Stephen D. Hursting (16). The cells and their subpassages were stored in the vapor phase of liquid nitrogen until use. All cells were visually monitored for obvious contaminants, such as bacterial and fungal growth. They were frequently tested for mycoplasma with a commercially available kit (MycoAlert® PLUS Mycoplasma Detection Kit). E0771 cells were cultured in DMEM media, 1 mM penicillin-streptomycin, 10% fetal bovine serum (FBS), glutamine, and 20mM HEPES at 37°C in 5% CO2 and 95% humidity. mWnt cells were cultured in DMEM media, 1mM penicillin-streptomycin, 10% FBS, glutamine, 5mM HEPES, 0.1% 2-mercaptoethanol at 37°C in 5% CO2 and 95% humidity.

### Gelatin Tumor Phantoms

Gelatin was dissolved in 100mL boiling deionized water with stirring. The solution solidified at room temperature overnight in dome-shaped silicone molds for tumor retention visualization and in cylindrical silicone molds for compression testing. For retention visualization, 50 or 100μL of food dye or 50 or 100μL of XCSgel_slow_ containing blue food dye was injected into a gelatin tumor phantom using a 26g needle.

### Compression Testing

Gelatin tumor phantoms were compressed using an Instron 5944 single-column mechanical testing system (Instron; Norwood, Massachusetts, USA) at a ramp rate of 0.20mm/s with a 500N load cell. Force (N) and displacement (mm) were recorded. Stress was calculated as force/cross-sectional area. Strain was calculated as displacement/initial length. Cross-sectional area and initial length were measured with digital calipers. The Young’s modulus was calculated as the slope of the stress-strain curve in the initial linear region before the inflection point.

### In vitro Cytokine Release

50μL of XCSgel_slow_ loaded with IFNα2 and IL-12 were prepared one day prior to release. Cytokines were released in 1mL of 10% FBS in 1xPBS. For each time point, all release media was removed to simulate sink conditions and stored at 4°C to prevent degradation. Protein concentrations were measured via ELISA.

### Cytokine Fluorophore Labeling

Cytokines IL-15 and IFNα2 were fluorescently labeled using Alexa Fluor™ 647 Microscale Protein Labeling Kit. IL-12 was labeled using Alexa Fluor™ 647 Protein Labeling Kit. The degrees of labeling, moles of fluorophore per mole of cytokine, were calculated to be 1.4 (IL-12), 0.6 (IL-15), 1.6 (IFNα2), according to manufacturer’s instructions.

### Tumor Cytokine Localization and Tissue Clearing

5×10^5^ E0771 cells were implanted in the 4^th^ mammary fat pad of C57BL/6 mice and allowed to grow until 199±47mm^3^. 50μL XCSgel_slow_ co-formulated with 5μg IL-12 labeled with AF647 (IL-12-AF647) or 5μg IL-12-AF647 in 50μL saline were delivered i.t. in E0771 tumors. Following delivery, tumors were excised and fixed in 1xPBS/4% paraformaldehyde at 4°C overnight. Tissues were cleared using the iDISCO clearing protocol (https://idisco.info/). Briefly, tumors were dehydrated with increasing concentrations (20-100%) of methanol at room temperature. Samples were then shaken with 66% dichloromethane/33% methanol at room temperature for three hours, followed by two fifteen-minute washes in 100% dichloromethane to remove excess methanol. Tissues were finally placed in dibenzyl ether for imaging. Cleared tumors were imaged using an LaVision UltraMicroscope II equipped with zoom body optics and an Olympus MVPLAPO 2X/0.5 NA lens with a corrected dipping cap. Zoom was set to 0.80x, resulting in a pixel size of 3.75µm. Z stacks were taken with 5µm spacing. The sample was illuminated with 3 sheets from one side (2 of them angled, to minimize shadowing), with the horizontal focus in the center of the field of a view, and a sheet NA setting of 0.016, which results in a 16.9µm sheet thickness. Fluorescence was collected using two channels: 488nm laser excitation and emission collection through a Chroma ET525/50 filter, for autofluorescence, and 647nm laser excitation and collection through a Chroma ET690/50 filter, for staining. Images were analyzed using IMARIS software version 10.1.1.

### In vivo Immunogenicity and Histology

5×10^5^ E0771 cells were implanted in the left 4^th^ mammary fat pad of C57BL/6 mice. When E0771 tumors reached 154±41mm^3^, mice were randomized into two groups, and 50μL of XCSgel_slow_ or XCSgel_fast_ was injected i.t.. On days 1, 3, and 7, mice were euthanized, and tissue collected. The tumor, gel, and surrounding tissue were dissected and fixed in 10% neutral buffered formalin. Tissue processing – including paraffin-embedding, sectioning, and staining – was performed by the Histopathology Laboratory at the NC State College of Veterinary Medicine. Blinded analysis of tissue reactions and immune cell infiltrates was performed by a board-certified veterinary pathologist (D.M.M.) on routine hematoxylin and eosin-stained slides.

### In vivo Fluorescence Imaging

For the variable cytokine release study, 50μL of XCSgel_slow_ loaded with 73.5nmol of AF647-labeled cytokine (IL-12, IL-15, or IFNα2) was injected subcutaneously on the shaved flank of female Balb/c mice. For the variable XCSgel with IL-12 study, 50μL of XCSgel_slow_ or XCSgel_fast_ loaded with 5μg (73.5nmol) of IL-12-AF647 was injected subcutaneously on the shaved flank of female Balb/c mice, with a saline medium as a control. Fluorescence (Ex/Em 640/690 nm) images were taken on a Spectral Instruments Imaging Ami HTX (Tucson, Arizona, USA) optical imaging system. Background values were subtracted, and the data were then normalized to the maximum efficiency in the experiment (17).

### Pharmacokinetics/Pharmacodynamics

5×10^5^ E0771 cells were implanted in the left 4^th^ mammary fat pad of C57BL/6 mice. Fourteen days after tumor implantation (tumor volumes 159±101 mm^3^), mice were randomized into three treatment groups. 50μL of XCSgel_slow_ co-formulated with 5μg IL-12 (XCSgel_slow_-IL-12_5μg_) or XCSgel_fast_-IL-12_5μg_ were delivered i.t., with a saline medium as a control. Separate cohorts of mice were bled one day prior to treatment and 4-, 24-, 48-, and 72-hours following treatment. Following each bleed, mice received 100-200μL injection of Hanks’ Balanced Salt Solution i.p. to restore lost liquid volume. Sera were collected and stored at −20°C until use. Serum IL-12 and IFNγ concentrations were determined via ELISA.

### Antitumor Studies

For the E0771 single orthotopic tumor study with variable cytokines, 5×10^5^ E0771 cells were implanted in the left 4^th^ mammary fat pad of C57BL/6 mice. When tumors reached 182±72mm^3^, (day 12 post implantation), mice were treated i.t. with 50μL of XCSgel_slow_-IL12_5μg_, XCSgel_slow_-IL15_30μg_, XCSgel_slow_-IFNα2_10μg_, XCSgel_slow_-GMCSF_5μg_, or left untreated. IL-15, IFNα2, and GM-CSF doses were determined using the maximum delivered dose in mice in preclinical studies, and the IL-12 dose was determined via prior work in our lab (18–20) (Supplemental Table 2). For the E0771 single orthotopic tumor study, 5×10^5^ E0771 cells were implanted in the left 4^th^ mammary fat pad of C57BL/6 mice. When tumors reached 125±57mm^3^, (day 17-21 post implantation), mice were treated i.t. with 50μL of XCSgel_slow_-IL12, XCSgel_slow_-IL12_1μg_, XCSgel_slow_-IL12_5μg_, or XCSgel_slow_-IL12_20μg_. Cured mice were rechallenged with 5×10^5^ E0771 cells on the opposite flank at least 100 days following initial treatment. Mice that rejected the E0771 rechallenge were rechallenged with 5×10^5^ mWnt cells on the left flank 95 days following the E0771 rechallenge. For the mWnt single orthotopic tumor study, 5×10^5^ mWnt cells were implanted in the left 4^th^ mammary fat pad of C57BL/6 mice. When tumors reached 114±63mm^3^, (day 9 post implantation), mice were treated i.t. with 50μL of XCSgel_slow_-IL12_5μg_ or were left untreated. For the abscopal E0771 tumor study, 5×10^5^ E0771 cells were implanted in the left 4^th^ mammary fat pad of C57BL/6 mice. To establish abscopal tumors, 5×10^5^ E0771 cells were implanted subcutaneously on the right flank three days after the primary tumor was implanted. When primary tumors reached 165±76 mm^3^ (16 days post implantation) and abscopal tumors were palpable, 50μL of XCSgel_slow_-IL12_5μg_ or XCSgel_fast_-IL12_5μg_ were delivered i.t., with XCSgel_slow_ and no treatment as controls. Tumor volumes were measured every 2-3 days and calculated using the formula: V = (w × w × l)/2, where V is tumor volume, w is tumor width and l is tumor length. Mice were euthanized when total tumor burden exceeded 2000mm^3^ for single tumors and 3000mm^3^ for two tumors, when tumors became ulcerated, or when mice became moribund.

### Spectral Flow Cytometry

Tissue samples were minced, incubated with Type IV collagenase and DNase I at 37°C for 30min with shaking at 250RPM, and then passed through 70-micron filters. Cell suspensions were then resuspended in phosphate buffered saline (PBS) with 1:800 Live/Dead Blue and incubated at room temperature for 10 minutes, protected from light in a 96-well V-bottom plate. Antibodies for extracellular antigens were prepared in Horizon^TM^ Brilliant staining buffer at a 2x dilution and added directly to the cell suspensions (Supplemental Table 3). Cells were stained for 30min at 4°C and protected from light. Stained cells were washed with PBS containing 2% fetal calf serum. Cells were then fixed and permeabilized, and intracellular antigens were stained using the eBioscience FoxP3/Transcription factor staining kit. Samples were run on a 5-laser Cytek Aurora following daily quality control experiments using SpectroFlow QC beads. Data were analyzed with FlowJo software (TreeStar, Version 10.9.0) and OMIQ (Dotmatics).

### Statistical Analysis

Sample size calculations were performed to detect effect sizes ranging from 1.6-2.2 among groups and to achieve 80% power at a significance level of 0.05. A standard two-tailed t test was used to test for significance between two sample groups. Differences between three or more treatment groups were determined using ANOVA with Tukey’s HSD post-hoc testing. The above analyses were conducted using Prism (GraphPad Software Inc., San Diego, California, USA).

## Results

### XCSgel effectively localizes cytokine immunotherapy in vivo when delivered intratumorally

Local retention of i.t. injections is limited by dense tumor architectures as well as high interstitial fluid and injection pressures, which exclude low viscosity injectates. To visualize the leakage of injectate, we developed transparent gelatin tumor phantoms that were mechanically similar to solid tumors. Compression testing of different concentrations of gelatin tumor phantoms revealed a Young’s modulus range of 9.8-41.9kPa (Supplemental Figure 1A). These data mapped onto the range of tumors’ Young’s Modulus, which for breast cancer, for example, range between 10-43 kPa (21). 14 and 10w/v% gelatin had Young’s moduli of 41.9±0.8 and 26.0±1.2 kPa, respectively. Blue dye in a saline solution injected into a gelatin tumor phantom offered a qualitative representation of poor retention, as most of the injectate returned via the needle track. (Figure 1A; Supplemental Movie 1A; Supplemental Figure 1B-C). When co-formulated with shear-thinning, self-healing XCSgel, the blue dye was completely retained within the gelatin tumor phantom (Figure 1B; Supplemental Movie 1B; Supplemental Figure 1B-C). These results were confirmed *in vivo* in the E0771 TNBC model. Fluorescently-labeled IL-12-AF647 in saline did not remain within the tumor when delivered *in vivo* and post-tissue clearing (Figure 1C), whereas IL-12-AF647 in XCSgel remained localized within the tumor when delivered following the same delivery and processing conditions (Figure 1D). These data demonstrate that XCSgel overcomes a major barrier, i.e. poor intratumoral retention, to the success of i.t. immunotherapy.

**Figure 1.**
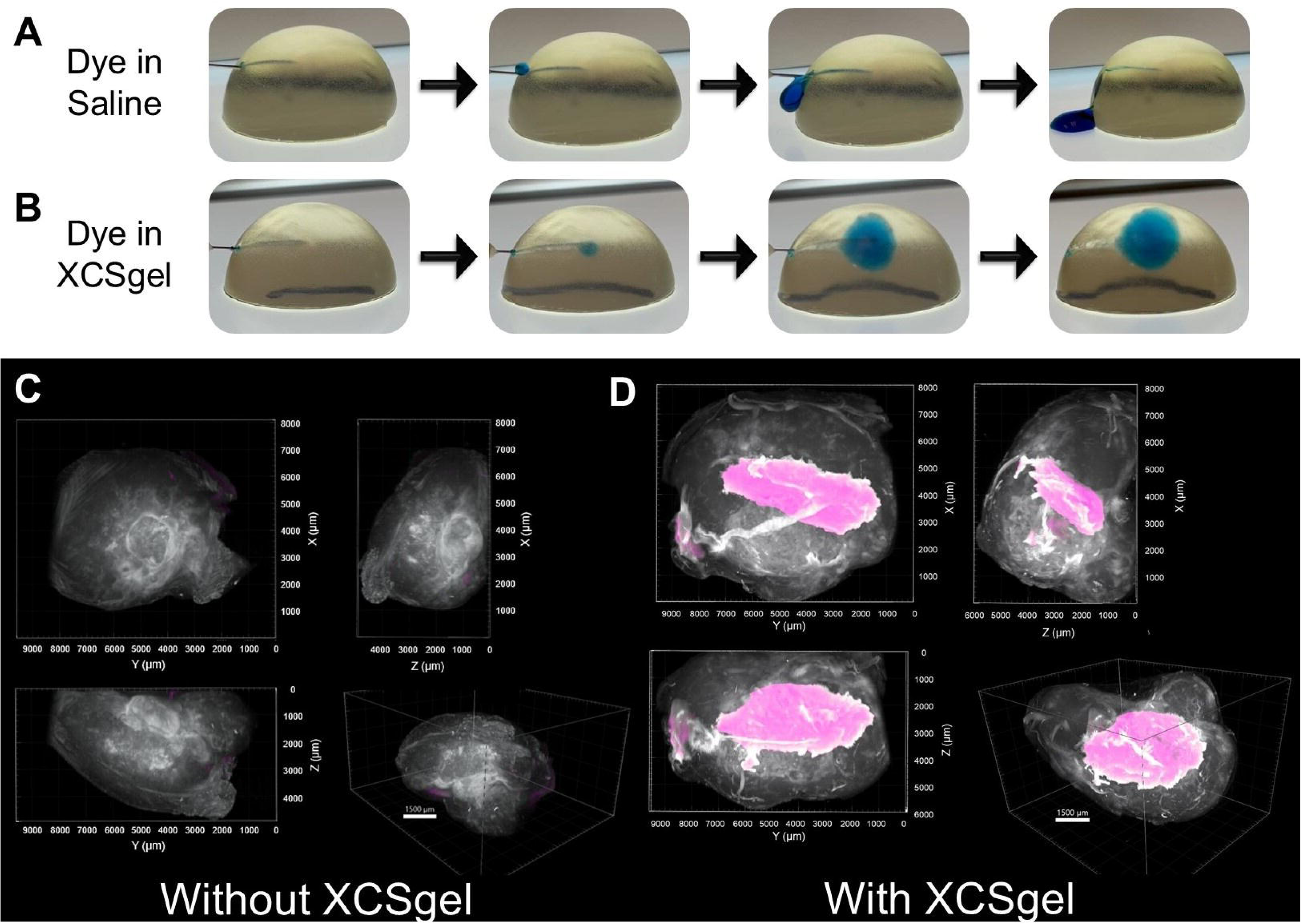
XCSgel as a delivery medium localizes therapies following i.t. delivery. (A) Dye in a saline solution was excluded following injection into gelatin tumor phantoms, while (B) dye within XCSgel_slow_ was retained within the gelatin tumor phantom. (C) *ex vivo* fluorescence imaging of i.t. delivered IL-12-AF647 within saline, which was not retained within a murine E0771 tumor *ex vivo*. (D) *ex vivo* fluorescence imaging of i.t. delivered IL-12-AF647, which was localized within a murine E0771 tumor when delivered in XCSgel_slow_.

### XCSgel alone induces a transient, tunable inflammatory response within injected tumors

Any foreign material delivered within the body will induce inflammation. The intensity and duration of which will vary based on the injected tissue, the host immune response, and the material delivered. In general, naturally derived materials, such as chitosan, elicit less inflammation compared to their synthetic counterparts. Knowledge of the type of immune response induced by a biomaterial can facilitate rational combinations of immunotherapies to leverage this response. To this end, the i.t. delivery of different compositions of XCSgel were delivered i.t. in the E0771 TNBC model. Two compositions, called XCSgel_slow_ and XCSgel_fast_ due their relative release kinetics, were chosen. XCSgel_slow_, had a high degree of deacetylation (DDA) (95%) and therefore a higher positive charge by virtue of its higher density of primary amine groups (Figure 2A). The second, XCSgel_fast_, had a reduced negative charge due to its 70% deacetylation and was co-formulated with a more negatively charged derivative of chitosan, CM-CS (Figure 2B). XCSgel_fast_ was previously found to release protein therapeutics more quickly *in vitro* with reduced inflammatory responses *in vivo* compared to XCSgel_slow_ (9). This trend agrees with literature finding higher inflammatory responses to positively charged versus negatively charged polymers (22–24).

**Figure 2.**
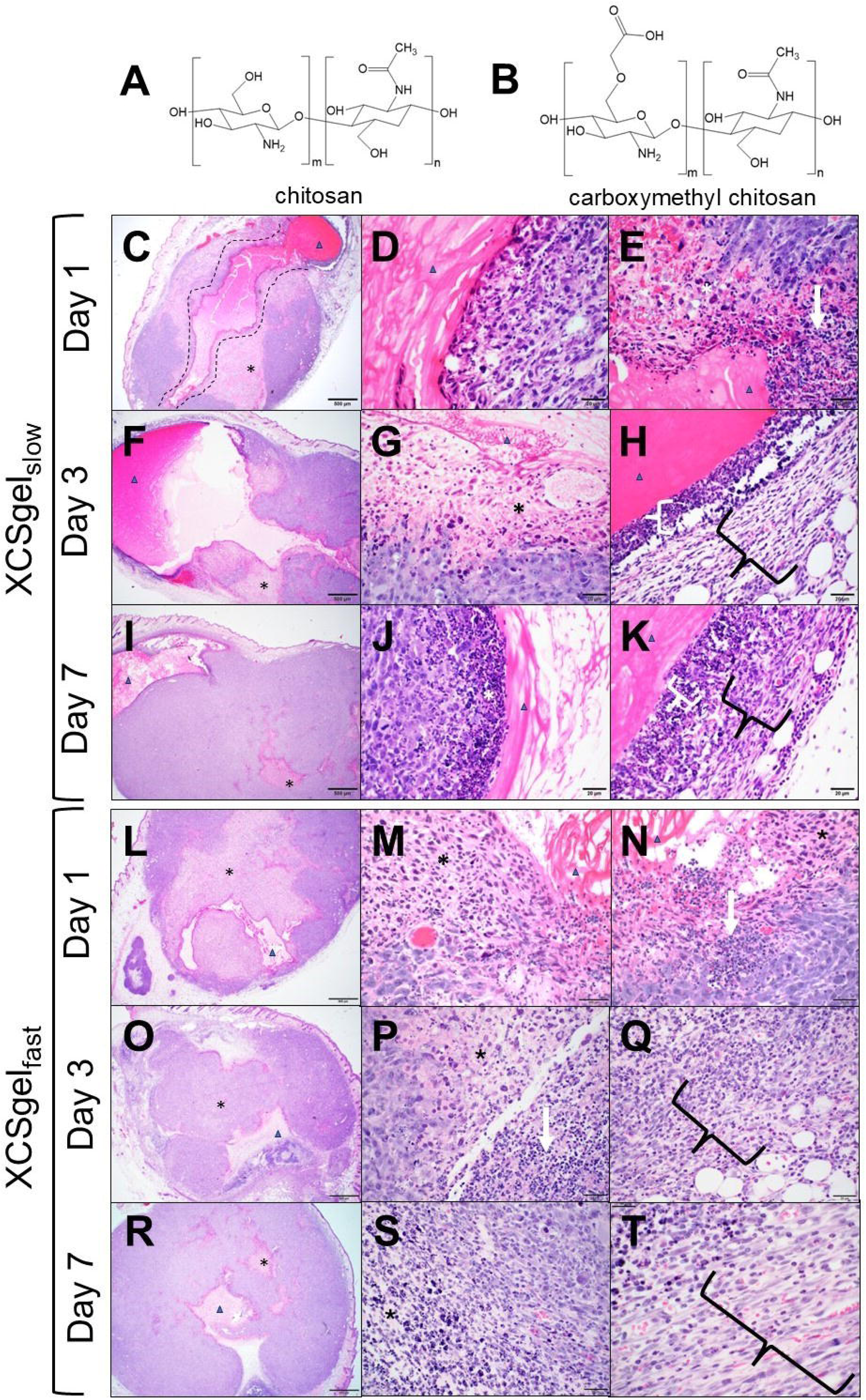
Histologic changes via H&E stain noted with i.t. injected XCSgel at different time points demonstrated inflammation that reduced over time. Chemical structures of (A) chitosan and (B) carboxymethyl chitosan. Two gels, (C-K) XCSgel_slow_ and (L-T) XCSgel_fast_, were evaluated. (C-E) XCSgel_slow_ day one. (C) Low magnification shows tissue track (dashed line) containing gel (arrowhead) from implant procedure with surrounding tumor necrosis (asterix). 2x objective. (D) Gel (arrowhead) and tumor interface showing early tumor necrosis (white asterix). 40x objective. (E) Gel (arrowhead) and tumor interface with tumor necrosis (white asterix) and marked neutrophilic inflammation (white arrow). 40x objective. (F-H) XCSgel_slow_ day three. (F) Low magnification shows persistent tissue track containing gel (arrowhead) with reduced tumor necrosis (asterix). 2x objective. (G) Gel (arrowhead) and tumor interface showing tumor necrosis (asterix). 40x objective. (H) Gel (arrowhead) with surrounding moderate neutrophilic inflammation (white bracket) and peripheral granulation tissue (black bracket). 40x objective. (I-K) XCSgel_slow_ day seven. (I) Low magnification shows tissue track containing gel (arrowhead) from implant procedure with minimal tumor necrosis (asterix). 2x objective. (J) Gel (arrowhead) and tumor interface showing a band of cell debris mixed with neutrophils (white asterix) but adjacent tumor necrosis is absent. 40x objective. (K) Gel (arrowhead) with surrounding mild neutrophilic inflammation (white bracket) and peripheral fibrosis (black bracket). 40x objective. (L-N)) XCSgel_fast_ day one. (L) Low magnification shows tissue track containing gel (arrowhead) from implant procedure with surrounding tumor necrosis (asterix). 2x objective. (M) Gel (arrowhead) and tumor interface showing early tumor necrosis (asterix). 40x objective. (N) Gel (arrowhead) and tumor interface with tumor necrosis asterix) and marked neutrophilic inflammation (white arrow). 40x objective. (O-Q)) XCSgel_fast_ day three (O-Q). (O) Low magnification shows persistent tissue track containing gel (arrowhead) with reduced tumor necrosis (asterix). 2x objective. (P) Tumor and tissue track interface showing on-going tumor necrosis (asterix) and less severe neutrophilic inflammation (white arrow). 40x objective. (Q) Tumor periphery with moderate neutrophilic inflammation and peripheral granulation tissue (black bracket). 40x objective. (R-T)) XCSgel_fast_ day seven. (R) Low magnification shows tissue track containing gel (arrowhead) from implant procedure with minimal tumor necrosis (asterix). 2x objective. (S) Tumor and tissue track interface showing a band of cell debris (necrosis) mixed with neutrophils (asterix). 40x objective. (T) Tumor periphery showing mild neutrophilic inflammation and peripheral fibrosis (black bracket). 40x objective.

Mice bearing E0771 tumors were treated i.t. with either gel, and timepoints were taken at day 1, 3, and 7 post-injection (Supplemental Table 4). One day post-injection, XCSgel_slow_ induced moderate i.t. inflammation, with 42±18% tumor necrosis (Figure 2C-E). Inflammation was neutrophilic, with polymorphonuclear leukocytes (PMNs) found around the gel and scattered lymphoplasmacytic (LP) inflammation around the periphery (Figure 2E). On day three, inflammation remained moderate, with 35±9% tumor necrosis (Figure 2F-H). Inflammation was neutrophilic and eosinophilic, with granulation tissue surrounding the gel and PMN inflammation. The granulation contained scattered LP inflammation (Figure 2H). One week post-injection, the severity of the inflammation had decreased to mild, with reduced necrosis within the tumor (Figure 2I-K). Inflammation remained both neutrophilic and eosinophilic, though it had noticeably decreased around the gel (Figure 2K). The presence of granulation tissue was increased at this timepoint with some progressing towards mature fibrous tissue.

XCSgel_fast_ induced mild to marked severity of inflammation after one day post-injection into the E0771 tumor. The gel induced tumor necrosis (57±6%), and inflammation was primarily neutrophilic (Figure 2L-N). On the third day post-injection, inflammation was moderate, with 28±6% tumor necrosis (Figure 2O-Q). Inflammation was both neutrophilic and eosinophilic (Figure 2P-Q). Gel was not visualized at this timepoint, but granulation tissue was observed as well as PMN infiltration in necrotic tumor regions. Seven days post-injection, inflammation was noticeably reduced to mild or minimal (Figure 2R-T). Tumor necrosis was noted at 30±9%. XCSgel was likely fully resolved as it could not be visualized, and PMN infiltration was observed in areas of tumor necrosis (Figure 2T).

It is important to note that the tumor necrosis induced by XCSgel alone was limited to the region directly in contact with the gel and did not result in tumor control, as demonstrated by a subsequent study (Figure 5C). Over time, XCSgels were resolved following i.t. injections in a similar manner as subcutaneous (s.c.) injections in prior studies (9).

### XCSgel releases cytokine therapeutics irrespective of charge or size

Cytokine immunotherapies can induce robust antitumor responses but are limited due to their high toxicity *in vivo* (3,25), making them prime candidates for i.t. delivery (4,27,28). Previously, XCSgel was found to tunably release protein therapeutics in a charge dependent manner *in vitro,* which underscores the importance of electrostatic interactions between charged polysaccharides and proteins (9). Therefore, to explore release kinetics in vivo, three clinically relevant antitumor cytokines, IL-12 (26), IL-15 (27), and IFNα2 (28), were chosen based on their distinct sizes, charges at neutral pH, and isoelectric points. These parameters are listed as follows: IL-12 (68kDa, −10.9 at pH 7.4, pI∼6.3), IL-15 (13kDa, −10.7 at pH 7.4, pI∼4.5), and IFNα2 (19kDa, +1.8 at pH 7.4, pI∼7.9) (29). The release profile of IFNα2 and IL-12 *in vitro* demonstrated significantly different release kinetics with the smaller, positively charged IFNα2 releasing much more quicky (Supplemental Figure 2). For *in vivo* studies, each cytokine was labeled with AF467 and delivered s.c. in XCSgel_slow_. All three cytokines displayed similar slow-release profiles and were imageable up to 28 days post-injection, with statistical significance only between IL-12 and IFNα2 at day 21 (Figure 3A-B). Otherwise, unlike our previous *in vitro* data (9), no difference between the cytokine release rates was observed, suggesting other parameters, such as the complex environment or the inflammatory tissue response *in vivo*, trumped the importance of electrostatic interactions.

**Figure 3.**
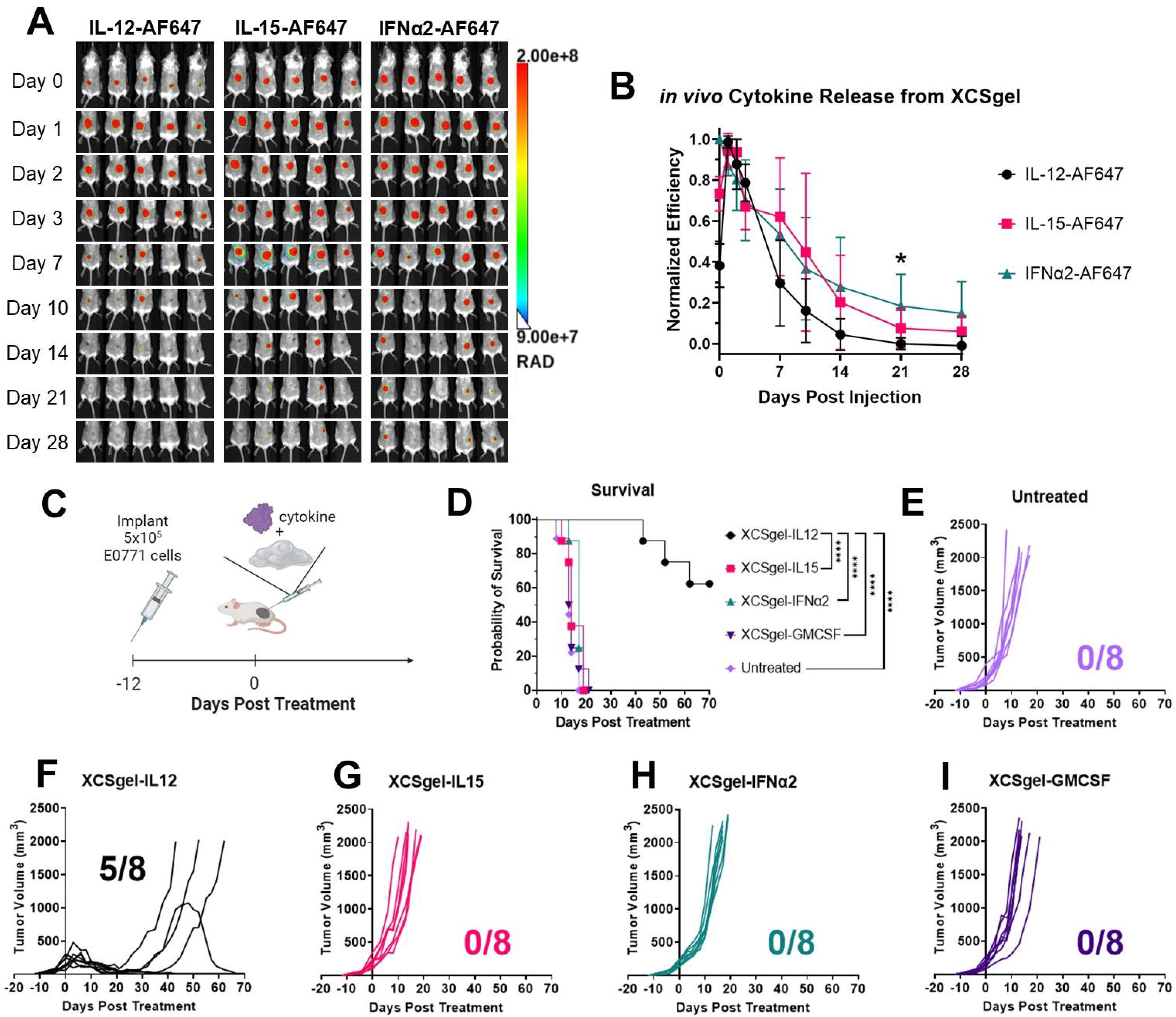
*In vivo* fluorescence imaging of labeled cytokines in XCSgels demonstrated long-term release, and anti-tumor studies display clear preference for IL-12. (A-B) Variable cytokine release. AF647-labled cytokines IL-12, IL-15, and IFNα2 were loaded into a different XCSgel_slow_ gels and delivered subcutaneously on the flanks of Balb/c mice. (A) Images at 640 nm/690 nm (Ex/Em) were taken on days 0, 1, 2, 3, 7, 10, 14, 21, and 28 post-injection. (B) Normalized efficiency values were taken and plotted over time. The significance was determined using one-way ANOVA and Tukey’s HSD posthoc testing. For IL-12-AF647 vs IFNα2-AF647, *P<0.01. n= 5 mice per experimental group. (C-I) Anti-tumor studies with variable cytokines. (C) Orthotopic primary E0771 tumors were established 12 days prior to treatment. (D) Survival and tumor growth curves of (E) untreated, (F) XCSgel_slow_-IL12_5μg_, (G) XCSgel_slow_-IL15_30μg_, (H) XCSgel_slow_-IFNα2_10μg_, and (I) XCSgel_slow_-GMCSF_5μg_ were monitored for 70 days post treatment. For the survival data, statistical significance was calculated using the Log-rank (Mantel–Cox) test. n = 8 mice per experimental group.

### XCSgel-IL12 outperforms other commonly used antitumor cytokines co-formulated in XCSgel

To compare antitumor activities, a range of antitumor cytokines (IL-12, IL-15, IFNα2, GM-CSF) commonly pursued in clinical studies (3,25,30) were co-formulated in XCSgel_slow_ and injected i.t. in an orthotopic E0771 primary tumor (Figure 3C). XCSgel_slow_-IL12 induced complete tumor rejection in 5 of 8 treated mice (Figure 3D). Tumor elimination was durable for at least 70 days (Figure 3F). A statistically significant slight initial tumor growth delay was noted in XCSgel-IFNα2, but this effect waned over time (Figure 3H). There were no discernible antitumor effects among untreated, XCSgel_slow_-IL15, XCSgel_slow_-IFNα2, and XCSgel_slow_-GMCSF cohorts (Figure 3E,G-I). Due to XCSgel_slow_-IL12’s superior performance compared to the other treatments, the remaining studies focused on localization of IL-12.

### Different compositions of XCSgel release cytokine IL-12 at different rates

As XCSgel_slow_ did not release different cytokines at distinct rates, the composition of XCSgel was engineered to determine if a single cytokine could be released at variable rates. Based on IL-12’s demonstrated antitumor activity (Figure 3F) and our prior work, IL-12 was chosen as a model cytokine therapeutic (19,31,32). IL-12-AF647 was co-formulated with both XCSgel_slow_ and XCSgel_fast_ and injected subcutaneously on the flanks of mice *in vivo*, with IL-12 delivered in saline as a control (Figure 4A). After 24 hours post-injection IL-12 delivered in saline was nearly undetectable, with high levels of cytokine remaining in both XCSgels (Figure 4A-C). XCSgel_fast_ retained IL-12 for about 2 weeks post-injection (Figure 3B). XCSgel_slow_ continued to release IL-12 over 3 to 4 weeks, with detectable levels up to day 28 (Figure 3A-C). Beginning at timepoint day 3, statistical differences in IL-12 release were observed between the two gels, which was again observed at days 10, 17, and 21 post-injection (Figure 3C). These results supported prior *in vitro* studies (9) demonstrating that XCSgel can be tuned to manipulate the rate of delivery *in vivo*.

**Figure 4.**
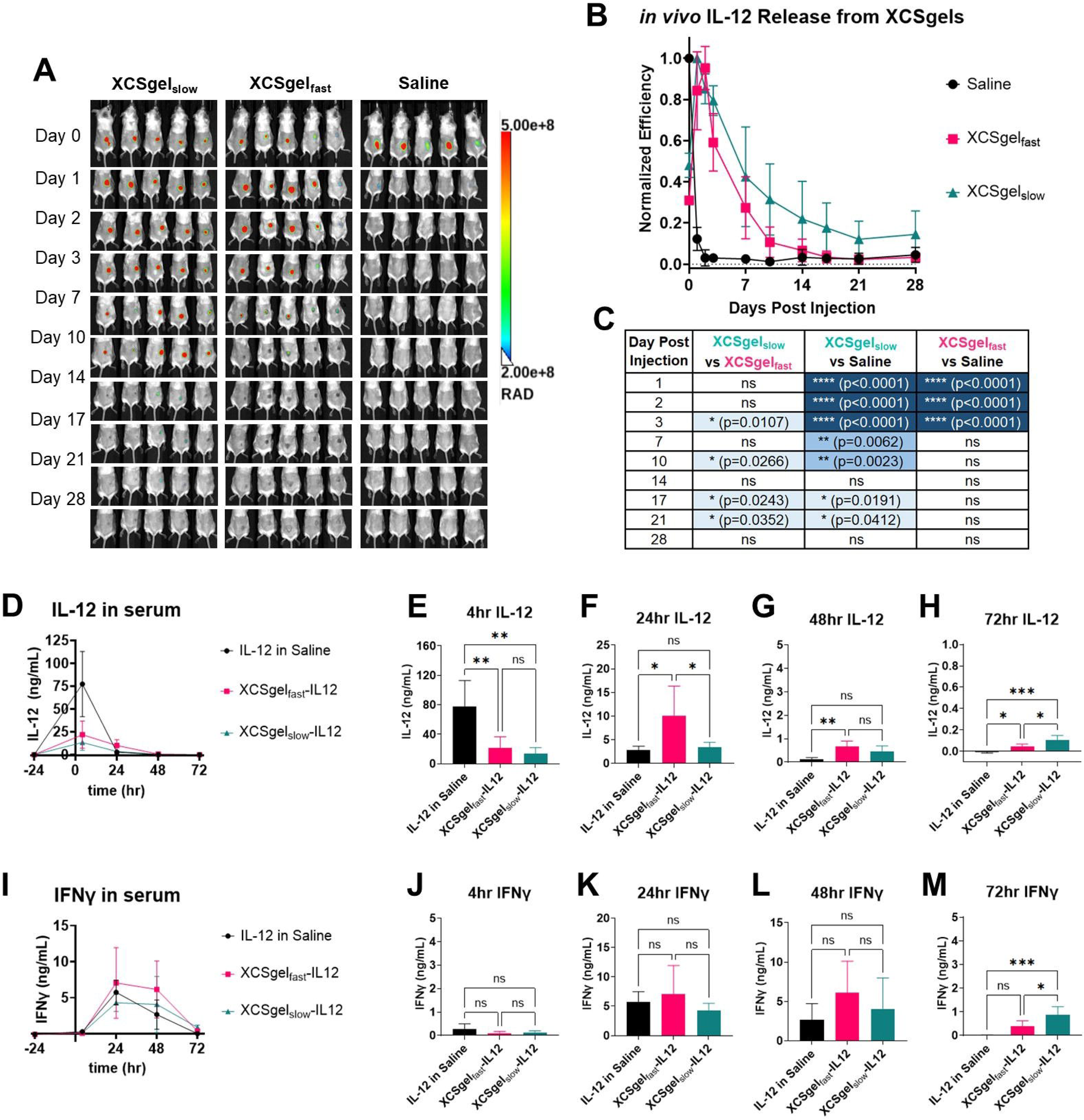
*In vivo* fluorescence imaging of IL-12 and pharmacokinetics of serum IL-12 and IFNγ following delivery of XCSgel-IL12 revealed tunable release. (A-C) Variable XCSgel release with IL-12-AF657. AF647-labeled IL-12 was loaded into XCSgel_fast_, XCSgel_slow_,, and saline and delivered subcutaneously on the flanks of Balb/c mice. (A) Images at 640 nm/690 nm (Ex/Em) were taken on days 0, 1, 2, 3, 7, 10, 14, 17, 21, and 28 post-injection. (B) Normalized efficiency values were taken and plotted over time. The significance was determined using one-way ANOVA and Tukey’s HSD posthoc testing and listed in (C). \**P* < 0.05, \*\**P* < 0.01, \*\*\*\**P* < 0.0001. n= 5 mice per experimental group. (D-M) Pharmacokinetics and pharmacodynamics of i.t. IL-12 in XCSgels. IL-12 (5 μg) was loaded XCSgel_fast_, XCSgel_slow_,, and saline and delivered intratumorally in orthotopic E0771 breast cancer tumors in C57BL/6 mice. Blood was collected from the facial mandibular vein at 4, 24, 48, and 72 hours post-injection. (D) IL-12 serum concentrations were plotted over time, with statistical significance plotted at (E) 4 hours, (F) 24 hours, (G) 48 hours, and (H) 72 hours. (I) IFNγ serum concentrations were plotted over time, with statistical significance plotted at (J) 4 hours, (K) 24 hours, (L) 48 hours, and (M) 72 hours. The significance was determined using one-way ANOVA and Tukey’s HSD posthoc testing. \**P* < 0.05, \*\**P* < 0.01, \*\*\**P* < 0.001, \*\*\*\**P* < 0.0001. n= 5 mice per experimental group.

### XCSgel reduces systemic IL-12 exposure

Systemic delivery of immunotherapies and cytokines leads to significant adverse events (3,25) and is thus dose limited. To determine if intratumoral delivery and retention of a potent cytokine, IL-12, within XCSgel mitigates systemic exposure, the pharmacokinetics of IL-12 were evaluated. XCSgel_slow_-IL12 and XCSgel_fast_-IL12 were injected i.t. in an E0771 TNBC orthotopic tumor. IL-12 alone served as a control. At timepoints 4, 24, 48, and 72 hours, mice were bled, and their sera collected for circulating cytokine concentrations of both IL-12 and IFNγ, a downstream cytokine produced by activated T and NK cells in response to IL-12 (33).

IL-12 alone caused a spike in serum IL-12 levels 4 hours post-i.t. injection, which then diminished significantly 24-hours post-i.t. injection with barely detectable levels at 48-hours and undetectable levels at 72-hours (Figure 4D). At 4 hours post-injection, XCSgel_fast_-IL12 treatment resulted in significantly lower serum IL-12 levels compared to unformulated IL-12 (Figure 4E). However, at 24-, 48-, and 72-hours, mice treated with i.t. XCSgel_fast_-IL12 had higher serum IL-12 concentrations than unformulated IL-12 although concentrations were quite low – less than 1ng/ml (Figure 4F-H). XCSgel_slow_-IL12 treatment resulted in the lowest levels of circulating IL-12 4 hours post-injection (Figure 4E). At 24, 48, and 72-hours post-injection, XCSgel_slow_-IL12 treatments maintained very low but measurable levels of IL-12 (Figure 4F-H). Taken together, these data demonstrated that co-formulation in XCSgel can reduce systemic exposure to IL-12 and that XCSgel can be modified to control systemic exposure. It is important to note that XCSgel markedly reduced but did not completely eliminate systemic exposure to IL-12. Whether this small amount of circulating IL-12 is beneficial to stimulating systemic antitumor immunity is the subject of ongoing studies.

Given that IL-12-induced toxicity is largely attributed to IFNγ (34), systemic circulation of this cytokine was also tracked. Circulating IFNγ levels peaked at 24 hours post injection for all IL-12 treatment groups (Figure 4I). IFNγ levels were reduced at 48 hours and still further reduced at 72 hours, indicating a resolution to baseline (Figure 4I). There were no statistical differences among treatment groups at the 4-, 24-, and 48-hour timepoints (Figure 4J-L) and only XCSgel_slow_ maintained a slightly elevated IFNγ concentration at 72-hours (Figure 4M).

### XCSgel-IL12 eliminates primary orthotopic TNBC tumors and induces protective immunity

The antitumor activity of i.t. XCSgel_slow_-IL12 at a range of IL-12 doses, 0, 1, 5, or 20µg, was evaluated in orthotopic E0771 tumors (Figure 5A). XCSgel-IL12_1µg_ treatment delayed tumor growth and extended survival compared to the XCSgel alone group (Figure 5B-D), however, none of the treated mice survived beyond 40 days. XCSgel-IL12_5µg_ treatment eliminated nearly all (6/7) tumors (Figure 5E). After complete regression, none of the treated tumors recurred, and mice remained tumor-free for 60 days (Figure 5B). Interestingly, increasing the dose of IL-12 in XCSgel to 20 µg reduced antitumor activity, with only 3/7 tumors completely regressing (Figure 5E). Cured mice from all groups were rechallenged in the opposite flank with E0771 cells and remained tumor-free, demonstrating protective immunity (Figure 5G). Following the E0771 rechallenge, mice were then challenged with the mWnt TNBC model. As expected, mWnt tumors grew in all mice, demonstrating immune specificity (Figure 5G).

**Figure 5.**
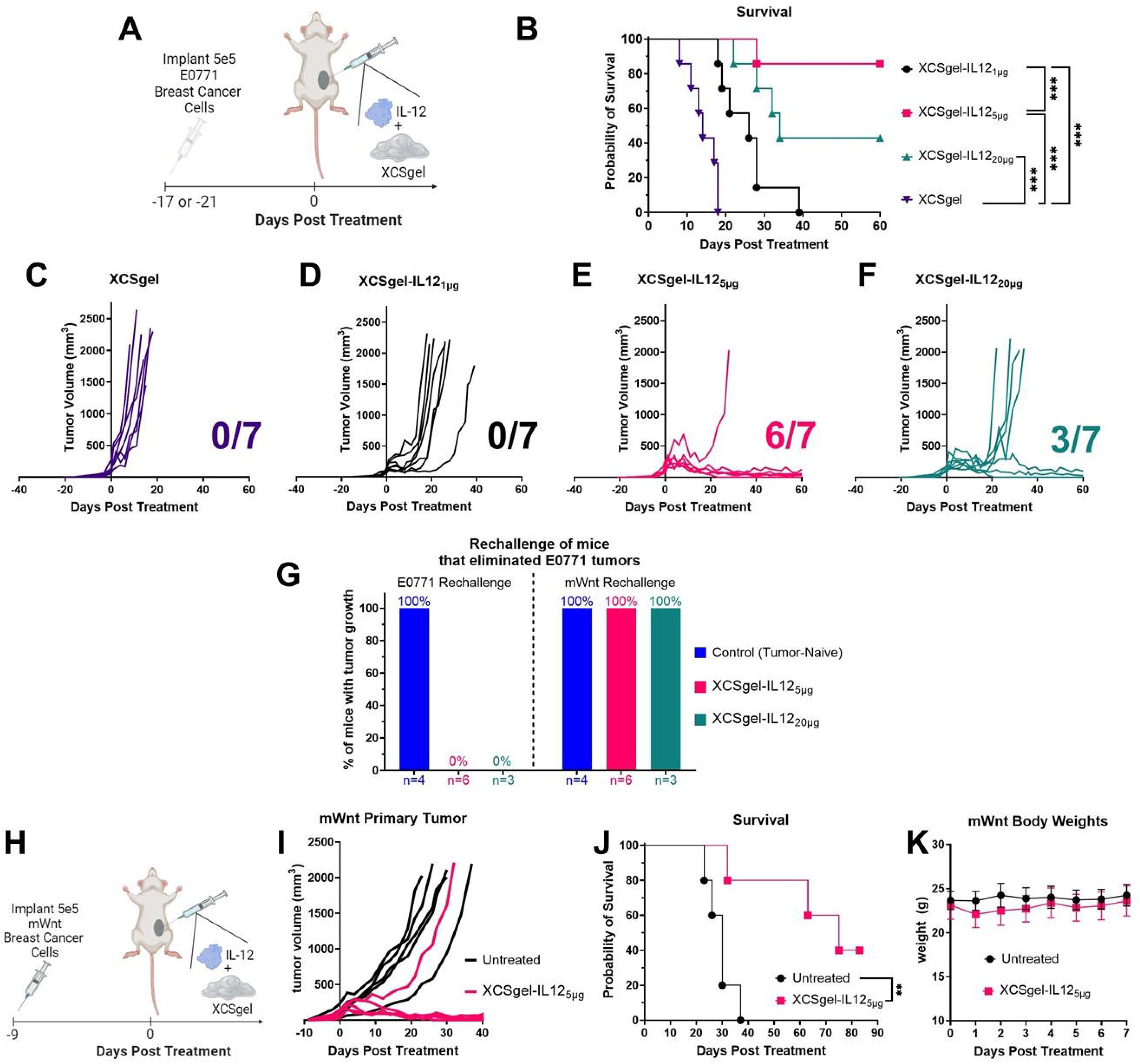
Intratumoral delivery of IL-12 co-formulated in XCSgel eliminated murine triple negative breast cancer tumors. (A-H) IL-12 was delivered within XCSgels_slow_ to orthotopic E0771 tumors at variable doses (0, 1, 5, 20 μg). (A) Orthotopic E0771 tumors were treated at either 17 or 21 days post tumor implantation with a single dose of IL-12 in XCSgel. (B) Survival and tumor growth curves of (C) XCSgel_slow_, (D) XCSgel_slow_-IL12_1μg_, (E) XCSgel_slow_-IL12_5μg_, (F) XCSgel_slow_-IL12_20μg_ were monitored for 60 days post treatment. For the survival data, statistical significance was calculated using the Log-rank (Mantel–Cox) test. (G) Mice whose tumors were eliminated were rechallenged with 5×10^5^ E0771 cells on the right flank, and their tumor growth was monitored. Mice that rejected the E0771 rechallenge were rechallenged with 5×10^5^ mWnt cells on the left flank. (H-K) IL-12 in XCSgel rejected TNBC mWnt tumors without toxicity. (H) Orthotopic mWnt tumors were treated nine days post tumor implantation with a single dose of XCSgel_slow_-IL12_5μg_. (I) mWnt tumor growth curves and (J) survival curves up to 85 days post treatment following treatment. (K) Body weights of treated mice, measured daily for seven days following treatment. The significance was determined using the Log-rank (Mantel-Cox) test. \*\*\**P* < 0.001. The number of mice per experimental group are as follows: (A-F) n = 7, (G) n = 5 additional naïve mice for E0771 rechallenge and n = 4 additional naïve mice for mWnt rechallenge, (H-K) n = 5.

To evaluate function in another, more difficult to treat tumor model, XCSgel_slow_-IL12_5µg_ was delivered i.t. in immunologically ‘cold’ mWnt orthotopic tumors (Figure 5I) (16). 1/5 mice experienced complete tumor rejection, with survival measured up to 85 days (Figure 5J). Body weight measurements were taken to assess potential systemic toxicities. however, no mouse experienced significant weight loss after XCSgel_slow_-IL12_5µg_ treatment (Figure 5K).

### XCSgel-localized IL-12 reshapes the T cell landscape within the TIME

To determine how XCSgel-IL12 alters the TIME, tumors were resected seven days after treatment when tumor volume differences became obvious. As illustrated in the UMAP plots in Figure 6A-C, high dimensional spectral flow cytometry revealed untreated and XCSgel-treated tumors displayed similar immune compositions, but XCSgel_slow_-IL12_5µg_ dramatically altered the composition of the TIME, particularly T cell populations (Figure 6A-C). Cell markers are listed in Supplemental Table 5.

**Figure 6.**
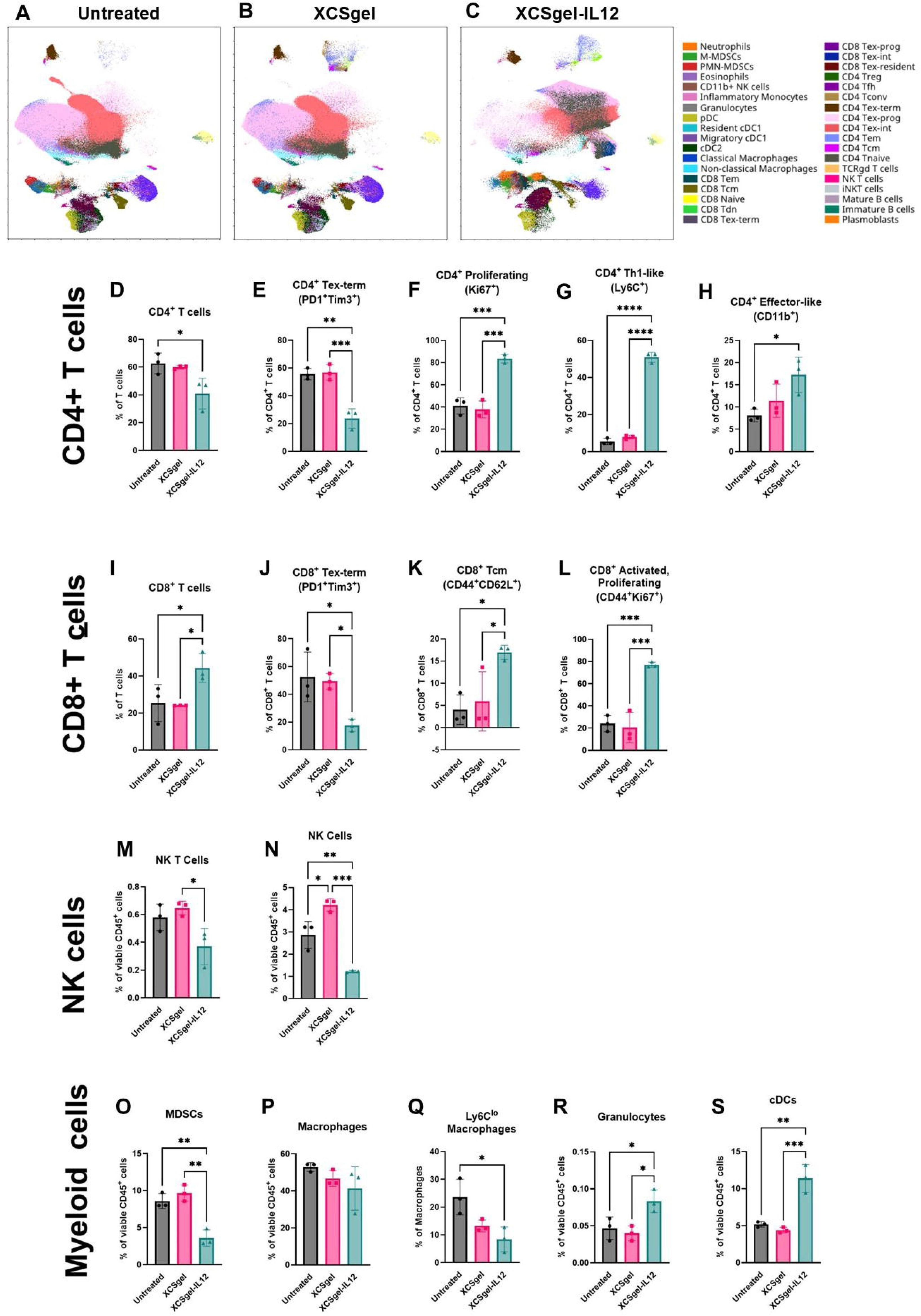
XCSgel-IL12 significantly alters the lymphoid cell compartment. Orthotopic primary E0771 tumors were established 12 days prior to treatment with XCSgel_slow_-IL12_5μg_ or XCSgel_slow._ Seven days after treatment when tumor volumes became palpable, tumors were resected and digested. Multi-spectral flow cytometry was run on tumor samples. (A-C) UMAP visualization of leukocytes populations and phenotypes of (A) untreated tumors, (B) tumors treated with or XCSgel_slow_,(C) tumors treated with XCSgel_slow_-IL12_5μg_. (D-X) Frequencies of cell populations and phenotypes. (D) CD4+ T cell population of T cells. For cell frequencies of CD4+ T cells, phenotypes include (E) terminally exhausted (PD-1+Tim3+), (F) proliferating (Ki67+), (G)Th1-like (Ly6C+), (H) effector-like (CD11b+). (I) CD8+ T cell population of T cells. For cell frequencies of CD8+ T cells, phenotypes include (J) terminally exhausted (PD-1+Tim3+), (K) central memory (CD44+CD62L+), (L) activated and proliferating (CD44+Ki67+). (M) NK T cell (CD3+NK1.1+) frequencies of CD45+ leukocytes, (P) NK cell (CD3-NK1.1+) frequencies of CD45+ leukocytes. Cells within the myeloid cell compartment include (O) myeloid derived suppressor cells, (P) macrophages, (Q) Ly6C^lo^ macrophages, (R) granulocytes, and (S) conventional dendritic cells. The significance was determined using one-way ANOVA and Tukey’s HSD posthoc testing. \**P* < 0.05, \*\**P* < 0.01, \*\*\**P* < 0.001, \*\*\*\**P* < 0.0001. n= 3 mice per experimental group.

The overall percentages of CD4^+^ T cells decreased from 63% to 41% of all T cells with XCSgel_slow_-IL12_5µg_ treatment (Figure 6D). Conversely, the frequency of CD8^+^ T cells increased from 25% to 44% with treatment (Figure 6I). More striking differences were found within these T cell lineages. The frequency of proliferating CD4^+^ T cells increased 2-fold, while the frequency of terminally exhausted CD4^+^ T cells decreased by 2.4-fold with XCSgel_slow_-IL12_5µg_ treatment (Figure 6E-F). A striking 9.4-fold increase in Ly6C^+^ Th1-like CD4^+^ cells was detected as well as a 2.1-fold increase in effector-like CD11b^+^ CD4^+^ cells (Figure 6G-H) (35,36). The frequency of terminally exhausted CD8^+^ T cells decreased by 3-fold while central memory-like CD8^+^ T cells, presumably containing favorable features of T cell stemness, increased by 4.2-fold (Figure 6J,K). Notably, the frequency of activated, proliferating CD8^+^ T cells increased by 3.2-fold with XCSgel_slow_-IL12_5µg_ treatment (Figure 6L). No differences were observed in the overall frequency of CD45^+^ leukocytes, T cells, regulatory T cells, and TCR γδ T cells (Supplemental Figure 3A-D). In contrast to beneficial changes in anti-tumor T cell populations, XCSgel_slow_-IL12_5µg_ decreased the frequency of NK T cells and NK cells by 1.6- and 2.3-fold, respectively (Figure 6M,N). Last, no dramatic differences in B cell populations were observed (Supplemental Figure 3E-H).

While the overall frequency of myeloid cells did not change with treatment, we detected changes in the composition of myeloid populations in the TIME (Supplemental Figure 3I). Myeloid derived suppressor cells (MDCSs) decreased 2.4-fold after XCSgel_slow_-IL12_5µg_ treatment, but the mononuclear (M-MDSCs) and polymorphonuclear (PMN-MDSCs) constituents exhibited no treatment-related changes (Figure 6O, Supplemental Figure 3J,K). Overall macrophage frequencies remained relatively stable; however, we detected a 3-fold decrease in the frequency of immunosuppressive Ly6C^lo^ macrophages, which have previously been shown to be pro-tumorigenic, in the XCSgel-IL12 group (Figure 6P,Q) (37). XCSgel_slow_-IL12_5µg_ also increased granulocytes from 0.04 to 0.08% and conventional dendritic cells (cDCs) over 2-fold from 5.2 to 11.4% (Figure 6R,S). No differences in neutrophils were noted (Supplemental Figure 3L).

### XCSgel-IL12 eliminates both primary orthotopic and abscopal flank TNBC tumors

A primary argument in favor of the use of systemic immunotherapies is for the treatment of disseminated disease. However, locally and rationally delivered immunotherapies can not only eliminate primary/treated tumors but also induce an adaptive immune response capable of eliminating untreated, distant tumors. To determine if XCSgel-IL12_5µg_ could eliminate abscopal tumors, a bilateral E0771 tumor model was developed in which the implantation of primary and abscopal tumors was staggered by three days. Sixteen days after primary tumor implantation, when abscopal tumors were palpable, XCSgel_slow_-IL12_5µg_ and XCSgel_fast_-IL12_5µg_ was injected i.t. in the primary tumor (Figure 7A). Controls included an untreated group and IL-12 alone. XCSgel_slow_-IL12_5µg_ led to long-term cures, with both primary and abscopal tumor elimination (Figure 7B-C). XCSgel_fast_-IL12_5µg_ treatment led to tumor rejection in 3/6 primary tumors, and only one mouse was cured of both the primary and untreated abscopal tumor (Figure 7D). There were no tumor regressions in either the IL-12 alone (Figure 7E) or untreated groups (Figure 7F). Comparisons of survival data between either XCSgel-IL12 group and the untreated cohort revealed statistically significant differences (Figure 7B). These data demonstrate that both injected and anenestic solid tumors can be eliminated with a single injection of XCSgel-IL12_5µg_.

**Figure 7.**
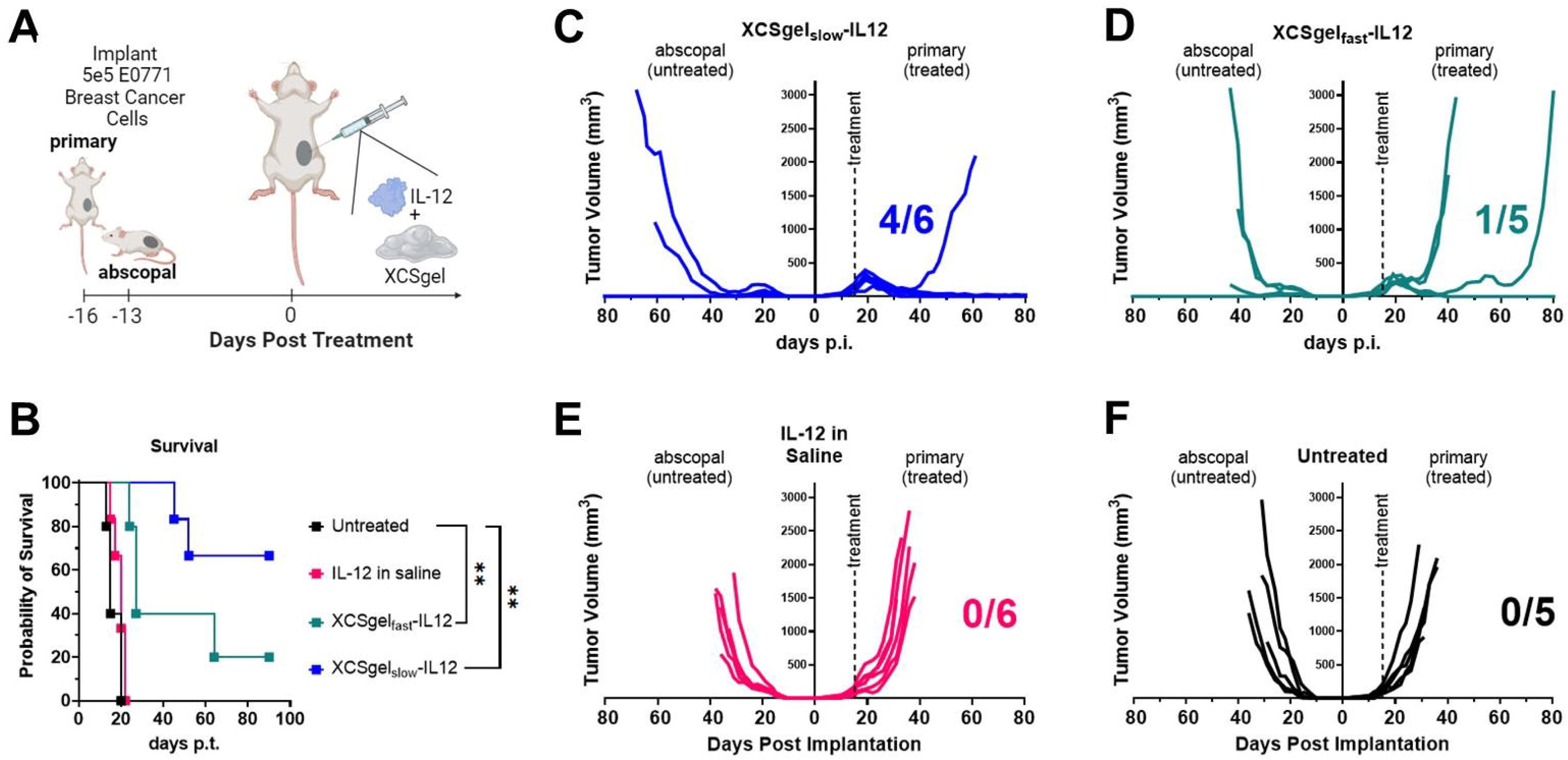
Intratumoral delivery of IL-12 co-formulated in XCSgel eliminated both primary and untreated abscopal E0771 tumors. (A) Orthotopic primary E0771 tumors were established 16 days prior to treatment, and untreated abscopal tumors were implanted on the flanks of mice 13 days prior to treatment. XCSgel-IL12_5µg_ was delivered i.t. into the orthotopic primary tumors. (B) Survival was monitored for 90 days post-treatment. Statistical significance was calculated using the Log-rank (Mantel–Cox) test. Tumor growth curves of the primary and abscopal tumors for (C) XCSgel_slow_-IL12_5µg_, (D) XCSgel_fast_-IL12_5µg_, (E) 5 μg IL-12 in saline i.t, and (F) no treatment were monitored for 80 days post-treatment. The significance was determined using the Log-rank (Mantel-Cox) test. \*\**P* < 0.01. n= 5-6 mice per experimental group.

## Discussion

I.t. immunotherapies promise to reduce irAEs while enhancing delivery to the tumor microenvironment. However, in the absence of a delivery system, i.t. injections are likely to rapidly leak from solid tumors, thus failing to maximize delivery to the tumor microenvironment. Of the 160 clinical trials currently investigating the i.t. delivery of immunotherapeutics, none utilized a delivery system, though different delivery strategies are in development, such as i.t. delivery of IL-12 linked to aluminum hydroxide (NCT06171750). I.t. injections of low viscosity solutions are not likely to be retained locally, as we have modeled *in vitro* and *in vivo* (Figure 1). To improve local retention, we previously developed XCSgel, an injectable, biodegradable, and clinically imageable hydrogel capable of coformulation and release of any number of immunobiologics (9). In stark contrast to saline-based injections, XCSgel effectively localizes its cargo within the tumors for sustained release (Figure 1).

XCSgel by itself induced a robust inflammatory response that was reduced over seven days after i.t. injection (Supplemental Table 4, Figure 2). The immune response was primarily neutrophilic at first, with eosinophils appearing 3 and 7 days after injection. Of particular note, XCSgel injected within a tumor induced a different reaction than XCSgel administered subcutaneously on the flanks of mice in prior data, which had both neutrophilic and granulomatous but not eosinophilic character (9). Location of implanted foreign materials have previously been demonstrated to impact the immune response (38). For example, fat necrosis often induces a more granulomatous response, which was seen with the subcutaneous injections. In comparing different XCSgel prototypes, the gel with the higher concentration, higher positive charge, and less hydrophobicity (XCSgel_slow_), was still present one week following injection, whereas the lower concentration, lower positive charge, and more hydrophobic, XCSgel_fast_ was not found one week following injection and induced less inflammation at later timepoints (Supplemental Table 4). This faster resolution of 70% deacetylated chitosan aligned with a prior study, which demonstrated that higher acetylation of chitosan led to faster degradation *in vivo* (39). Additionally, the incorporation of carboxylic acid groups via CM-CS may have also reduced inflammation, which draws the same conclusion as a prior study which demonstrated addition of carboxylic acid groups reduced the foreign body reaction (40). Both XCSgels induced tumor necrosis, which is typical of cationic polymers (41,42), and this necrosis was incapable of controlling the tumor (Figure 5C) and was reduced over time (Figure 2).

The immune reaction to XCSgel may have enhanced IL-12 activity and overall antitumor efficacy. Neutrophils produce IFNγ in response to IL-12 (43). Thus, the brisk XCSgel-induced neutrophilic infiltration likely enhanced the activity of co-formulated IL-12 in antitumor studies. In addition, the necrosis induced by XCSgel may provide a source of tumor antigen for the induction of an *in situ* vaccine response. Finally, DDA may have impacted antitumor efficacy. For example, a recent study delivering a highly deacetylated polymer derived from chitin demonstrated that a higher DDA led to a more robust antitumor response (44). Mechanistically, higher DDA led to higher induced mitochondrial stress, cGAS-STING activation, and IFNAR-dependent dendritic cell maturation (45).Taken together, XCSgel likely pairs well with potent pro-inflammatory immunotherapeutics, such as IL-12. Conversely, XCSgel is likely less effective and counterproductive when delivering anti-inflammatory therapeutics.

Our previous *in vitro* release studies demonstrated charge-dependent release of macromolecules from XCSgel, a phenomenon that was substantiated by prior research demonstrating the importance of charge on protein and polysaccharide interactions (9,46–48). To evaluate the impact of this charge-based release *in vivo*, three different cytokines with varying isoelectric points were formulated in XCSgel and delivered subcutaneously. Surprisingly, no difference in the release among the cytokines could be observed, though differences were observed *in vitro* (Figure 3A-B, Supplemental Figure 1). These findings suggest that the complex, proteinaceous *in vivo* environment may disrupt or shield any charge effects or that immune cells recruited by XCSgel-induced tissue responses may alter cytokine availability.

Nevertheless, additional *in vivo* imaging studies demonstrated that the release of a cytokine could be manipulated by changing the composition of XCSgel. Specifically, XCSgel_fast_ and XCSgel_slow_ were shown capable of maintaining measurable IL-12 in a tumor for 2 and 4 weeks, respectively (Figure 4A-B). Ongoing studies are exploring additional XCSgel compositions to further tune the release of diverse immunotherapeutics.

In addition to increasing localization of IL-12, XCSgel significantly reduced systemic exposure to IL-12 (Figure 4D-H). That XCSgel did not completely eliminate IL-12 dissemination may be a feature, as some amount of systemic IL-12, as long as it does not induce toxicity, may be beneficial. Serum IFNγ levels were in the low ng/ml range and remained largely unaffected, except at 72-hours post-treatment (Figure 4I-M), indicating that the relationship between IL-12 and IFNγ may not be proportional. Even though IL-12 related toxicity is attributed to IFNγ, the low levels measured did not result in observable clinical symptoms or significant body weight changes (Figure 5K). Most importantly, the enhanced local retention of IL-12 allows for reduced treatment frequency and likely a reduced number of clinic visits. It is useful to note that all antitumor experiments were performed with a single injection. In contract, t the severe IL-12-related toxicities discovered in initial clinical studies from 1997-2004 were a result of multiple IL-12 treatments (49).

Another important finding was the loss of antitumor activity when the dose of co-formulated IL-12 was increased from 5µg to 20µg (Figure 5E,F). These data indicate that the optimal biological dose of XCSgel-IL12 may not be the same as the maximum tolerated dose, a concept that has been advanced in immunotherapy circles (50). Because of the discrepancy between optimal and tolerated doses, the development or discovery of one or more biomarker(s) of for optimal biologic dose will be critical to the success of i.t. immunotherapies.

Immunoprofiling studies revealed that XCSgel_slow_-IL12_5μg_ induced profound changes in the TIME that likely contributed to the robust antitumor efficacy. Of note, XCSgel_slow_-IL12_5μg_ increased the frequency of activated and proliferating CD8^+^ T cells and reduced terminally exhausted CD4^+^ and CD8^+^ T cells, while reducing suppressive MDSCs (Figure 6). Similar changes would not be expected seven days after a single i.t. injection of IL-12 in saline, which we have shown has limited antitumor activity (31,32). Likely, it is the persistent Th1 signaling due to the sustained presence of IL-12 that drove alterations in the TIME. Future experiments will map out TIME changes as a function of time after XCSgel-IL12 in diverse tumor models.

In the context of TNBC, local and distant recurrences cause the greatest number of BC-related deaths (51). Rechallenge studies demonstrated that XCSgel-IL12 induced antitumor immunity that rejected live tumor rechallenges in a tumor-specific manner (Figure 5) as well as anenestic tumors (Figure 7). To evaluate the effectiveness of IL-12 co-formulated with XCSgel, an abscopal tumor on the opposite side of the primary tumor was implanted. Following treatment of only the primary tumor, 4/6 mice in XCSgel_slow_-IL12_5μg_ eliminated both tumors, with long-term survival (Figure 7B-C). Interestingly, XCSgel_slow_-IL12_5μg_ eliminated both treated and untreated tumors in 4/6 mice whereas XCSgel_fast_-IL12_5μg_ only eliminated both tumors in 1/5 mice (Figure 7D). This draws attention to the importance of release timing and/or delivery composition.

## Conclusion

XCSgel is a safe and effective delivery platform that addresses limitations and barriers to i.t. delivery, through the retention and slow release of immunotherapeutics. XCSgel enhanced intratumoral exposure to and antitumor activity of IL-12. A single injection of XCSgel-IL-12 drastically altered the TIME and facilitated robust systemic antitumor immunity. Data gathered supports the continued development of XCSgel-IL12 as an i.t. immunotherapy for TNBC.

## Acknowledgements

This work was performed in part by the Microscopy Services Laboratory, Department of Pathology and Laboratory Medicine, supported in part by P30 CA016086 Cancer Center Core Support Grant to the UNC Lineberger Comprehensive Cancer Center. Research reported in this publication was supported in part by the North Carolina Biotech Center Institutional Support Grant 2016-IDG-1016. We are grateful for the help that Dr. Pablo Ariel provided with light sheet imaging and IMARIS training. Schematic images were created using BioRender.com.

## Abbreviations

aCTLA-4: anti-cytotoxic T lymphocyte antigen 4
ATCC: American Type Culture Collection
BC: breast cancer
CAR: chimeric antigen receptor
cDC: conventional dendritic cell
CM-CS: carboxymethyl chitosan
CS: chitosan
CTL: cytotoxic T lymphocyte
DDA: degree of deacetylation
DN T cell: double negative T cell
EDC: 1-Ethyl-3-(3-dimethylaminopropyl)carbodiimide
FBR: foreign body reaction
FBS: fetal bovine serum
FDA: United States Food and Drug Administration
GRAS: generally recognized as safe
IFN: interferon
IL: interleukin
ILC: innate lymphoid cell
irAE: immune-related adverse event
i.p.: intraperitoneal
i.t.: intratumoral
LP: lymphoplasmacytic
MDSC: myeloid derived suppressor cell
M-MDSC: monocytic myeloid derived suppressor cell
NHS: N-Hydroxysuccinimide
NK: natural killer
PBS: phosphate buffered saline
pDC: plasmacytoid dendritic cell
PMN: polymorphonuclear leukocyte
PMN-MDSC: polymorphonuclear myeloid derived suppressor cell
s.c.: subcutaneous
TAM: tumor associated macrophage
Tcm: central memory T cell
Tconv: conventional T cell
Tfh: follicular helper T cell
Th: helper T cell
Tem: effector memory T cell
Tex: exhausted T cell
Tex-int: intermediate exhausted T cell
Tex-prog: exhausted progenitor T cell
Tex-term: terminally exhausted T cell
TIME: tumor-immune microenvironment
Tnaive: naïve T cell
TNBC: triple negative breast cancer
Treg: regulatory T cell

