## Supplemental Material for "An injectable chitosan hydrogel localizes and tunably releases immunotherapeutics intratumorally eliminating both treated and abscopal murine triple negative breast tumors"

**Supplemental Materials**

**Supplemental Table 1**. Reagents and resources used, listed along with their source and identifying information.

| Reagent or Resource | Source | Identifier |
|---|---|---|
| chitosan, 70% deacetylated, 100cp inherent viscosity | Heppe Medical Chitosan (Halle, Germany) | cat. #24204 |
| chitosan, 95% deacetylated, 100cp inherent viscosity | Heppe Medical Chitosan (Halle, Germany) | cat. #24704 |
| dichloromethane | Sigma Aldrich (Burlington, Massachusetts, USA) | cat. #270997-1L |
| dibenzyl ether | Sigma Aldrich (Burlington, Massachusetts, USA) | cat. # 108014-1KG |
| N-hydroxysuccinimide (NHS) | Sigma Aldrich (Burlington, Massachusetts, USA) | cat. #804518 |
| sodium hydroxide | Sigma Aldrich (Burlington, Massachusetts, USA) | cat. #795429-500G |
| fetal calf serum | Sigma Aldrich (Burlington, Massachusetts, USA) | cat. #C8056 |
| 2-mercaptoethanol | VWR (Radnor, Pennsylvania, USA) | cat. #97064-880 |
| dimethyl sulfoxide (DMSO) | VWR (Radnor, Pennsylvania, USA) | cat. #BDH1115-1LP |
| N-(3-Dimethylaminopropyl)-N'-ethylcarbodiimide hydrochloride (EDC) | VWR (Radnor, Pennsylvania, USA) | cat. #10118-238 |
| Alexa Fluor™ 647 Microscale Protein Labeling Kit | Fisher Scientific (Waltham, Massachusetts, USA) | cat. # A30009 |
| Alexa Fluor™ 647 Protein Labeling Kit | Fisher Scientific (Waltham, Massachusetts, USA) | cat. #A20173 |
| methanol | Fisher Scientific (Waltham, Massachusetts, USA) | cat. #A412P-4 |
| carboxymethyl chitosan (CM-CS) | Santa Cruz Biotechnology (Santa Cruz, California, USA) | cat. #SC358091 |
| hydrochloric acid | Honeywell International Inc. (Muskegon, Michigan, USA) | cat. #2033-1L |
| recombinant murine interleukin-15 (IL-15) | PeproTech (Cranbury, New Jersey, USA) | cat. #210-15 |
| Live/Dead Fixable Blue Dead Cell Stain Kit | Invitrogen (Waltham, Massachusetts, USA) | cat. #L23105 |
| eBioscience™ Foxp3 / Transcription Factor Staining Buffer Set | Invitrogen (Waltham, Massachusetts, USA) | cat. #00-5523-00 |
| recombinant murine interferon alpha 2 (IFNα2) | Invivogen (San Diego, California, USA) | cat. #34831285 |
| mIL-12 ELISA | Invivogen (San Diego, California, USA) | cat. #BMS6004 |

| Reagent or Resource | Source | Identifier |
|---|---|---|
| mIL-15 ELISA | Invivogen<br>(San Diego, California, USA) | cat. #900K188 |
| mIFNα ELISA | Invivogen<br>(San Diego, California, USA) | cat. #BMS6027 |
| mIFNγ ELISA | Invivogen<br>(San Diego, California, USA) | cat. #88-7314-88 |
| MycoAlert® PLUS Mycoplasma Detection Kit | Lonza<br>(Walkersville, Maryland, USA) | cat. #LT07-703 |
| gelatin<br>(Knox® gelatine) | Winland Foods<br>(Oak Brook, Illinois, USA) | Amazon, ASIN #B001UOW7D8 |
| 96-well V-bottom plate | Corning<br>(Corning, New York, USA) | cat. #3897 |
| Dulbecco's Modified Eagle Medium (DMEM) | Corning<br>(Corning, New York, USA) | cat. #15-013-CV |
| glutamine | Corning<br>(Corning, New York, USA) | cat. #25-005-CI |
| HEPES | Corning<br>(Corning, New York, USA) | cat. #25060CI |
| penicillin-streptomycin | Genesee Scientific<br>(El Cajon, California, USA) | cat. #25-513 |
| fetal bovine serum (FBS) | Genesee Scientific<br>(El Cajon, California, USA) | cat. #25-550 |
| Horizon™ Brilliant staining buffer | BD<br>(Franklin Lakes, New Jersey. USA) | cat. #566349 |
| SpectroFlo® QC Beads | Cytek<br>(Fremont, California, USA) | cat. #B7-10001<br>Lot #2004 |

**Supplemental Table 2.** Maximum delivered doses in mice in preclinical studies, listed alongside the publication PMID.

| Cytokine | Maximum *in vivo* dose / mouse | PMID |
|---|---|---|
| IL-15 | 2.5 µg | 19383782 |
| IL-15 | 2.5 µg | 20581314 |
| IL-15 | 3 ug | 32461349 |
| IL-15 | 5 ug | 20924130 |
| IL-15 | 5 ug | 37461129 |
| IL-15 | 10 ug | 15630141 |
| IL-15 | 10 µg | 19359475 |
| IL-15 | 28.8 µg | 19723883 |
| IL-15 | 30 µg | 7553894 |
| IFNα | 500 ng | 33976143 |
| IFNα | 1,250 units | 35441749 |
| IFNα | 6,400 units | 18813777 |
| IFNα | 10,000 units | 31541153 |
| IFNα | 10,000 units | 25520086 |
| IFNα | 50,000 units | 20309938 |
| IFNα | 64,000 units | 16953837 |
| IFNα | 100,000 units | 24550491 |
| IFNα | 100,000 units | 7584666 |
| GM-CSF | 100 ng | 19223554 |
| GM-CSF | 100 ng | 25701675 |
| GM-CSF | 650 ng | 21076828 |
| GM-CSF | 1 µg | 34605223 |
| GM-CSF | 1.5 µg | 17082611 |
| GM-CSF | 1.5 µg | 26265369 |
| GM-CSF | 2 ug | 17568998 |
| GM-CSF | 2 µg | 12759428 |
| GM-CSF | 5 µg | 38543209 |

**Supplemental Table 3**. Antibodies used for multi-spectral flow cytometry.

| Marker | Channel | Marker | Clone | Fluorophore | Supplier | Cat. # | Purpose | Dilution (ng) |
|---|---|---|---|---|---|---|---|---|
| 1 | UV2 | CD45 | 30-F11 | BUV395 | BD | 564279 | Leukocytes | 66.7 |
| 2 | UV6 | Viability | 16-10A1 | LIVE DEAD Blue | Invitrogen | L34961 | live/dead | 1.2 |
| 3 | UV9 | CD44 | IM7 | BUV536 | BD | 741227 | T cell activation (EM/CM) | 50 |
| 4 | UV11 | CD80 | 16-10A1 | BUV661 | BD | 741515 | M1 marker | 100 |
| 5 | UV14 | CD11c | HL3 | BUV737 | BD | 564986 | DC marker | 100 |
| 6 | UV16 | CD4 | RM4-4 | BUV805 | BD | 741913 | CD4 lineage | 100 |
| 7 | V3 | Ki67 | SolA15 | eF450 | Invitrogen | 48-5698-82 | Proliferating cells | 66.7 |
| 8 | V5 | TIM3 | 5D12 | BV480 | BD | 747618 | T cell exhaustion (Tex) | 100 |
| 9 | V7 | Ly6G | 1A8 | BV510 | BioLegend | 127633 | Neutrophils | 133.3 |
| 10 | V8 | CD62L | MEL-14 | BV570 | BioLegend | 104433 | L-selectin | 100 |
| 11 | V10 | CD115 | AFS98 | BV605 | BioLegend | 135517 | Neutrophils vs MDSCs | 100 |
| 12 | V11 | CD185 (CXCR5) | L138D7 | BV650 | Biolegend | 145517 | Tfh cells | 100 |
| 13 | V13 | CD69 | H1.2F3 | BV711 | Biolegend | 104537 | Activation (Trm) | 133.3 |
| 14 | V14 | CD138 | 281-2 | BV750 | BD | 747070 | Plasma cells (B cells) | 100 |
| 15 | V15 | F4/80 | BM8 | BV785 | Biolegend | 123141 | Dendritic Cell lineage | 100 |
| 16 | B2 | CD127 | A7R34 | AF488 | BioLegend | 135017 | ILC's & MP CD8s | 250 |
| 17 | B3 | IA/IE | M114.1 | Spark Blue 550 | BioLegend | 107661 | MHCII (B cells, myeloid cells, APCs) | 83.3 |
| 18 | B9 | CD11b | M1/70 | PerCP-Cy5.5 | BioLegend | 101228 | myeloid lineage cells | 66.7 |
| 19 | B10 | KLRG1 | 2F1 | PerCP-eFluor710 | Invitrogen | 46-5893-82 | Teff cells | 66.7 |
| 20 | YG1 | TCRγδ | GL3 | PE | BioLegend | 118108 | gamma delta T cells | 100 |
| 21 | YG3 | CD8a | 53-6.7 | Alexa Fluor 594 | BioLegend | 100758 | CD8 lineage | 250 |
| 22 | YG5 | NK1.1 | PK136 | PE/Cy5 | Biolegend | 108716 | NK cell lineage | 100 |
| 23 | YG6 | CD19 | eBio1D3 | PE/Cy5.5 | Thermofisher | 35-0193-82 | B cell lineage | 50 |
| 24 | YG9 | CD103 | 2 E7 | PE-Cy7 | BioLegend | 121426 | Trm & DC marker | 100 |
| 25 | YG10 | B220 | RA3-6B2 | PE-fire810 | BioLegend | 103287 | B cell & myeloid marker | 50 |
| 26 | R1 | FoxP3 | FJK-16s | APC | Thermofisher | 17-5773-82 | T regulatory cells | 100 |
| 27 | R2 | SiglecF | E50-2440 | AF647 | BD | 562680 | Eosinophils | 100 |

| Marker | Channel | Marker | Clone | Fluorophore | Supplier | Cat. # | Purpose | Dilution (ng) |
|---|---|---|---|---|---|---|---|---|
| 28 | R4 | CD3 | 17A2 | Alexa Fluor 700 | BioLegend | 100216 | T cell receptor | 250 |
| 29 | R7 | PD-1 | 29F.1A12 | APC-Cy7 | BioLegend | 135224 | T cell exhaustion (Tex) | 100 |
| 30 | R8 | Ly6C | HK1.4 | APC/Fire810 | BioLegend | 128055 | Myeloid cells | 66.7 |

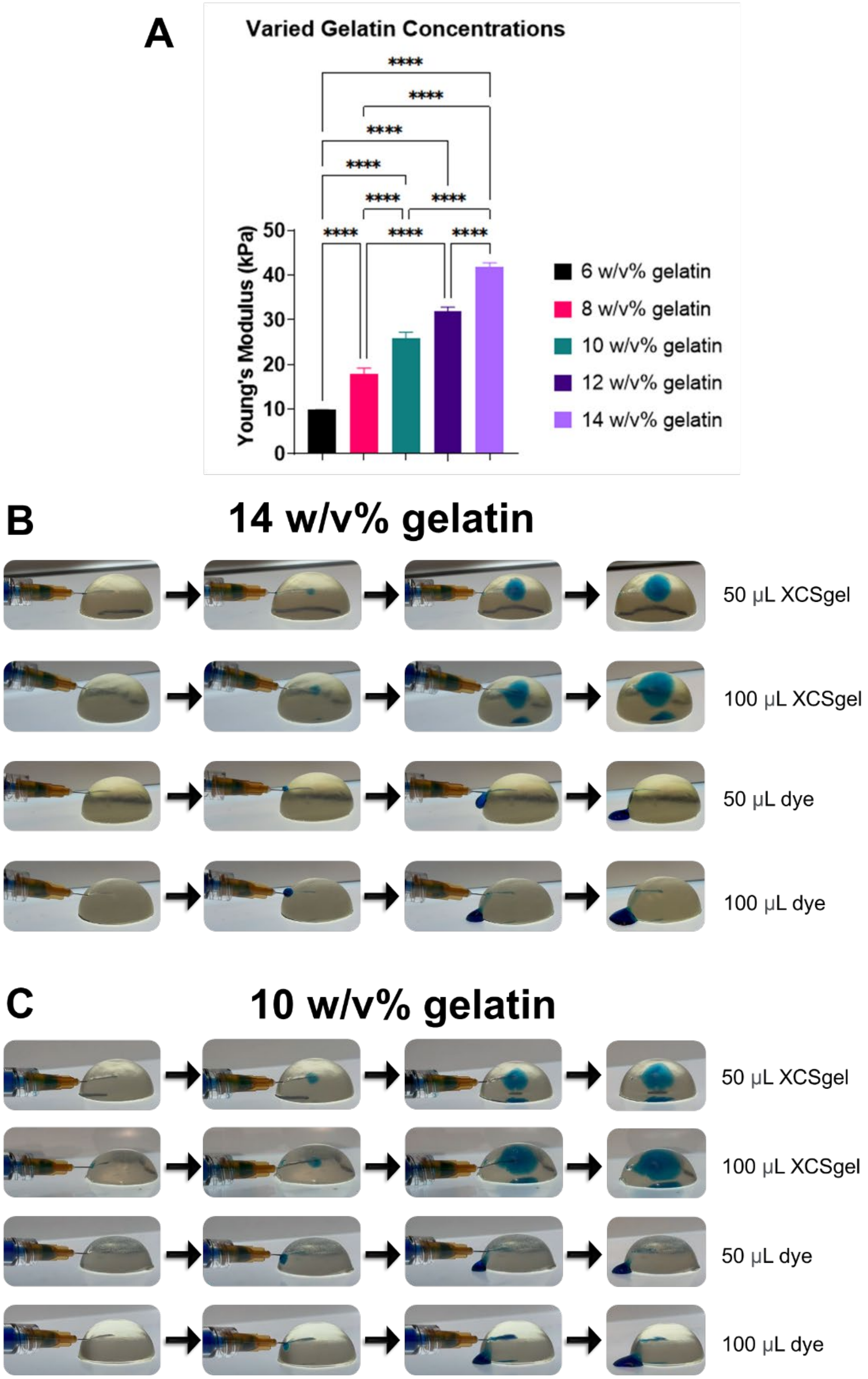


**Supplemental Figure 1**. Retention testing of XCSgel and saline in gelatin tumor phantoms. (A) Young's moduli of gelatin tumor phantoms at various gelatin concentrations. (B) Injection of 50μL XCSgel-dye, 100μL XCSgel-dye, 50μL dye in saline, and 100μL dye in saline in 14 w/v% gelatin. (C) Injection of 50μL XCSgel-dye, 100μL XCSgel-dye, 50μL dye in saline, and 100μL dye in saline in 10 w/v% gelatin.

**Supplemental Table 4**. Histologic changes noted with intratumorally injected XCSgel at different time points.

| XCSgel | Timepoint (day) | Inflammation (Y/N) | Severity | Necrosis | Type |
|---|---|---|---|---|---|
| $XCSgel_{slow}$ | 1 | Y | Moderate | Yes, tumor | Neutrophilic |
| | 3 | Y | Moderate | Yes, tumor | Neutrophilic and eosinophilic |
| | 7 | Y | Mild | Yes, tumor | Neutrophilic and eosinophilic |
| $XCSgel_{fast}$ | 1 | Y | Mild to Marked | Yes, tumor and around gel | Neutrophilic; one also with scattered macrophages |
| | 3 | Y | Moderate | Yes, tumor | Neutrophilic; two also eosinophilic |
| | 7 | Y | Minimal to Mild | Yes, tumor | Neutrophilic; one also eosinophilic |

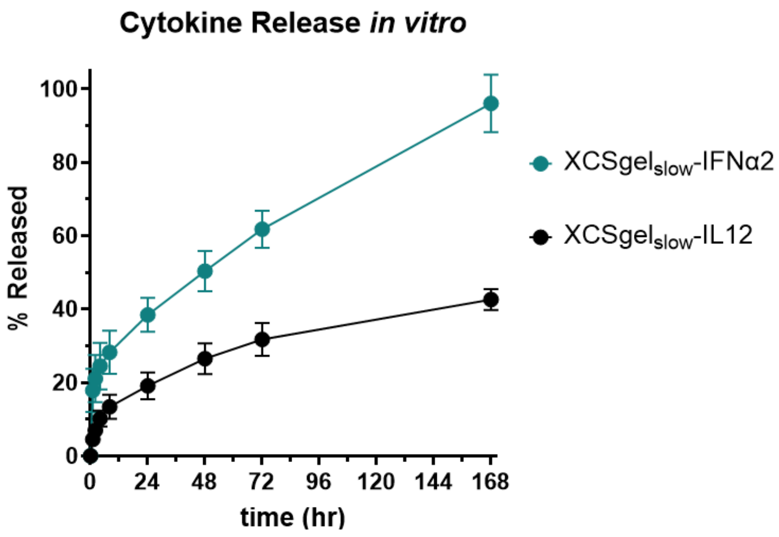


**Supplemental Figure 2.** *In vitro* release of cytokines IFNα2 and IL-12 from XCSgel$_{slow}$ over a period of 7 days.

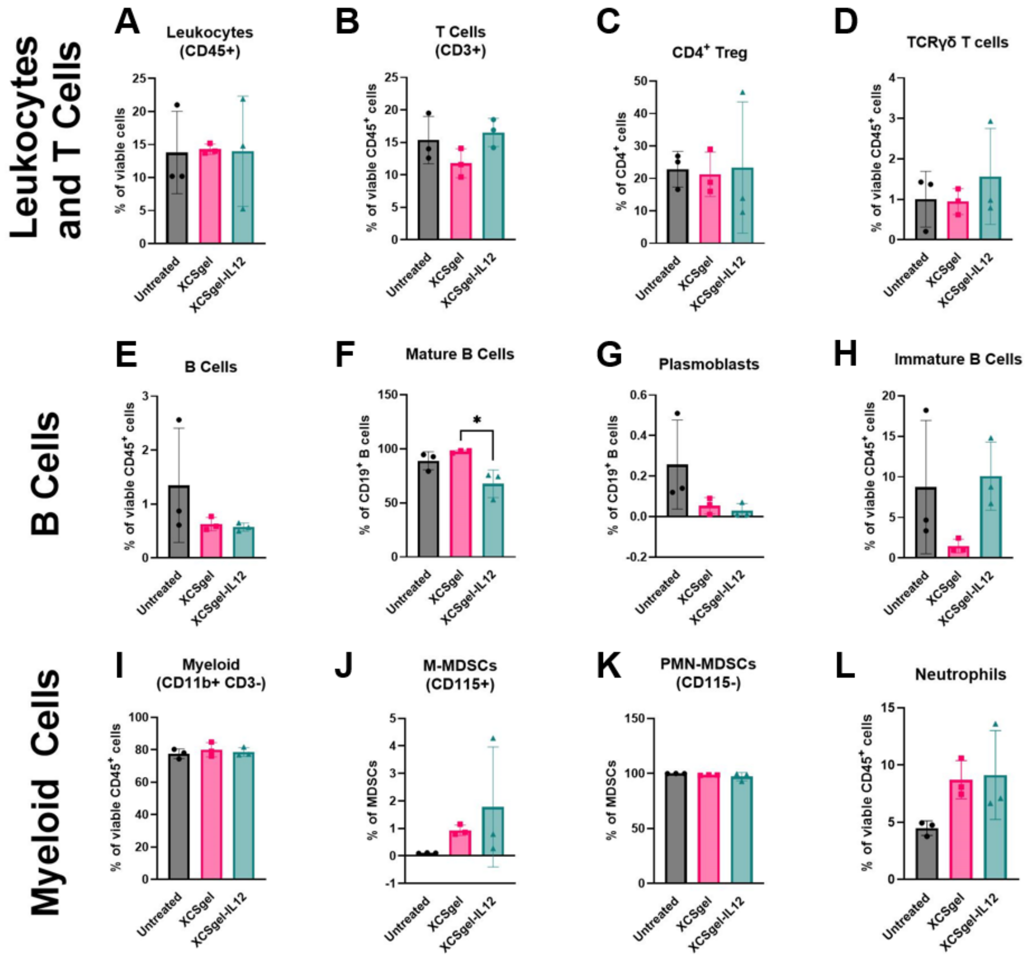


**Supplemental Figure 3.** Additional cell populations and phenotype frequencies following XCSgel-IL12 treatment. Orthotopic primary E0771 tumors were established 12 days prior to treatment with XCSgelslow-IL125µg or XCSgelslow. Seven days after treatment when tumor volumes became palpable, tumors were resected and digested. Multi-spectral flow cytometry was run on tumor samples. (A) CD45+ leukocyte frequencies of all viable cells. (B) $CD3^+$ T cell frequencies of all viable $CD45^+$ leukocytes. (C) $FoxP3^+$ regulatory T cell frequencies of $CD4^+$ T cells. (D) $TCR\gamma\delta^+$ T cell frequencies of all $CD4^+$ T cells. (E) Overall $CD19^+$ B cell frequencies of all viable $CD45^+$ leukocytes. (F) $B220^+CD138^-$ mature B cell, (G) $CD138^+B220^+$ plasmoblast, and (H) $CD138^-B220^-$ immature B cell frequencies of all $CD19^+$ B cells. (I) Myeloid cell frequencies of all viable $CD45^+$ leukocytes. (J) Myeloid and (K) polymorphonuclear MDSC frequencies of all MSDCs. (L) Neutrophil frequencies of all viable $CD45^+$ leukocytes. The significance was determined using one-way ANOVA and Tukey's HSD posthoc testing. $^*P < 0.05$. n= 3 mice per experimental group.

**Supplemental Table 5**. Markers used in flow cytometry to identify cell populations, with their respective abbreviations.

| Cell Type/Phenotype | Markers |
|---|---|
| T cells | Live CD45+CD3+ |
| $CD4^+$ T cells | Live CD45+CD3+CD4+ |
| Conventional $CD4^+$ T cell (Tconv) | Live CD45+CD3+CD4+CXCR5-FoxP3- |
| $CD4^+$ Bystander | Live CD45+CD3+CD4+CXCR5-FoxP3-PD1- |
| Central Memory $CD4^+$ T cell (Tcm) | Live CD45+CD3+CD4+CXCR5-FoxP3-PD1-Tim3+CD62L+ |
| Effector Memory $CD4^+$ T cell (Tem) | Live CD45+CD3+CD4+CXCR5-FoxP3-PD1-Tim3-CD62L- |
| Naïve $CD4^+$ T cell (Tnaive) | Live CD45+CD3+CD4+CXCR5-FoxP3-PD1-Tim3-CD62L- |
| Intermediate Exhausted $CD4^+$ T cell (Tex-int) | Live CD45+CD3+CD4+CXCR5-FoxP3-PD1+CD44+Tim3-Ly6C- |
| Exhausted progenitor $CD4^+$ T cell (Tex-prog) | Live CD45+CD3+CD4+CXCR5-FoxP3-PD1+CD44+Tim3-Ly6C+ |
| Terminally Exhausted $CD4^+$ T cell (Tex-term) | Live CD45+CD3+CD4+CXCR5-FoxP3-PD1+CD44+Tim3+Ly6C- |
| Follicular Helper $CD4^+$ T cell (Tfh) | Live CD45+CD3+CD4+CXCR5+ |
| Regulatory T cell (Treg) | Live CD45+CD3+CD4+FoxP3+ |
| Double Negative (DN) T cells | Live CD45+CD3+CD4-CD8- |
| $CD8^+$ T cells | Live CD45+CD3+CD8+ |
| $CD8^+$ Bystander | Live CD45+CD3+CD8+PD1- |
| Central Memory $CD8^+$ T cell (Tcm) | Live CD45+CD3+CD8+PD1-CD62L+CD44+ |
| Double Negative $CD8^+$ T cell (Tdn) | Live CD45+CD3+CD8+PD1-CD62L-CD44- |
| Effector Memory $CD8^+$ T cell (Tem) | Live CD45+CD3+CD8+PD1-CD62L-CD44+ |
| Naïve $CD8^+$ T cell (Tnaive) | Live CD45+CD3+CD8+PD1-CD62L+CD44- |
| Exhausted $CD8^+$ T cell (Tex) | Live CD45+CD3+CD8+PD1+ |
| Intermediate Exhausted $CD8^+$ T cell (Tex-int) | Live CD45+CD3+CD8+PD1+Ly6C-Tim3- |
| Exhausted Progenitor $CD8^+$ T cell (Tex-prog) | Live CD45+CD3+CD8+PD1+Ly6C+Tim3- |
| Exhausted Resident $CD8^+$ T (Tex-resident) | Live CD45+CD3+CD8+PD1+CD103+CD69+ |
| Terminally Exhausted $CD8^+$ T (Tex-term) | Live CD45+CD3+CD8+PD1+Ly6C-Tim3+ |
| Natural Killer (NK) T cells | Live CD45+CD3+NK1.1+ |
| $TCR\gamma\delta^+$ T cells | Live CD45+CD4+NK1.1-TCRgd+ |

| Cell Type/Phenotype | Markers |
|---|---|
| B cells | Live CD45+CD3-CD19+ |
| Immature B Cells | Live CD45+CD3-CD19+CD138-B220- |
| Mature B Cells | Live CD45+CD3-CD19+B220+ |
| Plasmoblasts | Live CD45+CD3-CD19+CD138+ |
| NK cells | Live CD45+CD3-CD11b+CD19-NK1.1+ |
| Myeloid | Live CD45+CD3-CD11b+CD19-NK1.1- |
| Monocytic (M)-MDSCs | Live CD45+CD3-CD11b+CD19-NK1.1-Ly6G+CD115+Ly6C- |
| Neutrophils | Live CD45+CD3-CD11b+CD19-NK1.1-Ly6G+CD115+Ly6C+ |
| Polymorphonuclear (PMN)-MDSCs | Live CD45+CD3-CD11b+CD19-NK1.1-Ly6G+CD115-Ly6C- |
| Eosinophils | Live CD45+CD3-CD11b+CD19-NK1.1-Ly6G-SiglecF+ |
| Macrophages | Live CD45+CD3-CD11b+CD19-NK1.1-Ly6G-SiglecF-F4.80+ |
| Conventional Dendritic Cells (cDCs) | Live CD45+CD3-CD11b+CD19-NK1.1-Ly6G-SiglecF-F4.80-MHCII+B220- |
| cDC2s | Live CD45+CD3-CD11b+CD19-NK1.1-Ly6G-SiglecF-F4.80-MHCII+B220-CD103-CD8a- |
| Migratory cDC1s | Live CD45+CD3-CD11b+CD19-NK1.1-Ly6G-SiglecF-F4.80-MHCII+B220-CD103+ |
| Resident cDC1s | Live CD45+CD3-CD11b+CD19-NK1.1-Ly6G-SiglecF-F4.80-MHCII+B220-CD8a+ |
| Plasmacytoid Dendritic Cells (pDCs) | Live CD45+CD3-CD11b+CD19-NK1.1-Ly6G-SiglecF-F4.80-MHCII+B220+ |
| Granulocytes | Live CD45+CD3-CD11b+CD19-NK1.1-Ly6G-SiglecF-F4.80-MHCII-Ly6C- |
| Inflammatory Monocytes | Live CD45+CD3-CD11b+CD19-NK1.1-Ly6G-SiglecF-F4.80-MHCII-Ly6C+ |
